# GENETIC VARIABILITY AND ASSOCIATION OF TRAITS IN COMMON BEAN (*Phaseolus vulgaris* L.) GENOTYPES IN SEKOTA, NORTH WESTERN ETHIOPIA

**DOI:** 10.1101/2024.08.21.609046

**Authors:** Abebe Assefa, Dereje Worku, Muluken Bantayhu, Fentaw Asres

**Affiliations:** Amhara Regional Agricultural Research Institute (ARARI), Sekota Dry Land Agricultural Research Centre (SDARC) P.O. Box, 62, Sekota, Ethiopia; Bahir Dar University College of Agriculture and Environmental Sciences (CAES) P.O. Box, 551, Bahir Dar, Ethiopia

**Keywords:** Cluster analysis, correlation analysis, heritability, principal component analysis

## Abstract

Common bean (Phaseolus vulgaris L.) is one of the most important pulse crops in Ethiopia, contributing to income generation and food security. Information on the genetic variability and trait associations of common bean in the Sekota district of north Western Ethiopia is inadequate. This study was initiated with the objective to assess variability, heritability and trait association among 64 common bean genotypes for quantitative traits using 8x8 simple lattice design at Aybra main research site 2023 under rain fed conditions. Analysis of variance was performed using SAS software and the ANOVA revealed highly significant variations among genotypes for all the traits considered in the study. The yield ranged from 1633.1 to 3702.10 kg ha^-1^ with a mean of 2542.53 kg ha^-1^. There was a yield advantage of 24.56 to 55.89% over the checks. A moderate genotypic coefficient of variation coupled with high heritability and high genetic advance as a percentage of the mean was obtained for plant height, branches per plant, hundred seed weight, seed yield, and harvest index. Branches per plant, aboveground biomass and harvest index had significant positive correlations and direct effects on seed yield at the genotypic and phenotypic levels while days to maturity had a significant negative correlation and indirect effect with seed yield at the genotypic. The maximum inter cluster distance was found between clusters VII and 8 (D^2^ =351.39), followed by clusters V and VIII (D^2^ =331.23). The first five principal component axes accounted for 74.3% of the total variation, with eigenvalues greater than unity. The number of days to maturity, plant height, number of pods per plant, number seeds per pod, seed yield, and harvest index were the traits that contributed most of the variation in the first PCs. Generally, the presence of variability, and strong positive association of traits among the genotypes were observed in the traits under study. Therefore, selection based on agronomic performance and hybridization based on cluster distance could be possible for the improvement of common bean in the study area.

## INTRODUCTION

Common bean (*Phaseolus vulgaris L*. 2n=22) popularly known as the dry bean, has massive, pinnately compound trifoliate leaves (Katungi *et al.,* 2009). According to FAOSTAT; (2019) the five top producer countries of dry beans in average annual are India (5.8 Mt), Myanmar (4.9 Mt), Brazil (3.0 Mt), the United States (1.3 Mt) and Mexico (1.2 Mt), followed by China and several African countries such as Tanzania, Uganda, Kenya and Ethiopia. For the past forty years, common beans have been one of Africa’s most important exportable pulse crops, with Ethiopia being the continent’s top exporter (Ferris and Kaganzi, 2008).

Common beans are widely grown by Ethiopian farmers due to their early maturity and ability to withstand terminal moisture deficit (Adem Mohammed and Estifanos Feleke, 2022). In Ethiopia mong the total area covered by pulse crops, white and red common beans constitute 103,288.55 and 208,295.03 hectares of land, respectively, with changes of 7.13% in the red type and 6.48% in the white type from 2020 to 2021 (CSA, 2021). In the Amhara region, the white common bean covers 53,504.27 hectares of land with a production of 900,961.3 qt and a productivity of 16.84 q ha^-1^ and the red type covers an area of 36073.59 hectare of land with the production of 683,189.14qt and the productivity of 18.94 q ha^-1^. In the Wag Hemira zone, 923.74 ha of land covered with red beans with a production of 6,462.68 qt and productivity of 7q ha^-1^ and 1,302.33 of white type covered areas, with a with production and productivity of 10,710.53 and 8.22 q ha^-1^, respectively (CSA, 2021). In Ethiopia, common bean is a significant national economic crop that contributes to the country’s food security. With 9.5% of all agricultural exports, it is Ethiopia’s third most valuable commodity and it is a staple food legume used in a variety of regional dishes across the nation. It is made and eaten in a variety of recipes, including kik, shiro, nifro, and sambosa and it can be used with other grains to make soup (Berhane Amsalu *et al.,* 2017) and use as medicine, animal feed, and honeybee foraging of flowers, helps to ensure food and nutrition security, generates revenue for smallholder farmers, and increases Ethiopia’s foreign exchange revenues in both domestic and international marketssuch as relay, double, and intercropping. It significantly contributes to soil fertility by fixing atmospheric nitrogen and preserving the stability and diversity of agricultural systems Mukankusi (2003), Demelash Bassa (2018), and Fitsum Alemayehu (2022).

To create new varieties, breeders must first understand genetic variability and the associations between traits of interest (Scossiroli *et al.,* 1963; Begum *et al.,* 2016). The effectiveness of a breeding program is mostly determined by the population’s level of genetic variability, the degree to which the targeted traits are strongly heritable, and the positive correlation and direct effect of the traits (Majumder *et al.,* 2008). Selecting the optimal genotypes based solely on grain yield can be difficult due to the complexity of grain production, which is a quantitative trait controlled by multiple genes and affected by both yield-related characteristics and environmental factors. In common bean, various studies have been conducted on genetic variability, diversity and traits association in different regions of Ethiopia and have reported the existence of variations and trait associations for various among tested genotypes/varieties (Kedir Gelgelu *et al*., 2019; Kedir Shafi *et al.,* 2019; Nigussie Kefelegn *et al.,* 2020; Masreshaw Yirga, *et al*., 2022, and Kedir Yimam *et al.,* 2023).

The development of suitable improved varieties of common bean through different breeding programs is one of the solutions to increase productivity and production. Such improvement of crops requires the generation and/or introduction of genetic variation and extensive evaluation of breeding materials at multiple locations (Negash Gelata *et. al.,* 2015). Therefore, for a successful breeding program, the presence of a sufficient amount of genetic variability/diversity in the population and the extent to which the desired traits are heritable, strong and positive direct correlation of traits play a vital role. Effectiveness of selection in any crop depends on the extent and nature of phenotypic and genotypic variability present in different agronomic traits. Such studies on common bean are insufficient, and there is no fully documented on genetic parameter or association information in the specified area (Wag-Hemira administrative zone). Genetic variability and heritability study in available common bean genotypes for morpho-agronomic traits is vital in common bean growing areas such as Wag-Hemria to develop new varieties and thereby increase yield and common bean has versatile importance and is grown in Wag-Hemira areas. However, its productivity in the area is very low (8.22 q ha^-1^) compared to the regional (17.89 q ha^-1^) and national averages (18 q ha^-1^) due to the use of unimproved seeds and improved common bean varieties under production is limited or almost null. The absence of improved common bean varieties coupled with biotic and abiotic factors are the major causes of the low productivity of common bean in Wag-Hemira areas. Therefore, precise information on the nature and magnitude of genetic variability and genetic relationships among common breeding materials is a prerequisite and is of per paramount important for common bean breeding.

Therefore this work was initiated with the objective of

To estimate genetic variability for quantitative traits among small seeded common bean genotypes;

To assess the association of yield and yield related traits

To identify the traits accounting for much of the total variation among the genotypes using multivariate analysis (PCA); and

To determine genetic similarity among genotypes using multivariate analysis

## MATERIALS AND METHODS

### Description of Experimental Location

The field experiment was conducted at the Sekota Dry Land Agricultural Research Centre, Aybra main research station, during the 2023 main cropping season. Aybra is located at 12^0^ 43^′^ 38″ N E longitude, 79^0^ 01′ 08″ E N latitude with an altitude of 1915 m.a.s.l. The site receives a mean minimum and maximum annual rainfall of 492 and 621 mm, with mean minimum and maximum temperatures of 15.4°C and 26.9°C, respectively Sekota Dry Land Agricultural Research Centre annual report (SDARC, 2016/17). The study site is located in the Sekota district, Wag Hemira Zone of the Amhara National Regional State (ANRS) (Figure 1), which is 448 kilometers (km) away from Bahir dar (the regional capital of Amhara) and 738 kilometres (km) away from Addis Ababa (capital of Ethiopia). The dominant soil type is classified as vertisol. The general slope at the site varies between 0 and 8% but can normally be found in the 0-25% slope range (Demlie Gebresellassie and Shawel Abebe, 2016). Sorghum (*Sorghum bicolor*), tef *(Eragrostis tef (zucc.)*) (Trotter), and barley (*Hordeum vulgare L*.) are the cereals, and common bean *(Phaseolus vulgaris L.),* mung bean (*Vigna radiata*), and filed pea (*Pisum sativum L*.) are some of the crops cultivated in the area (district) (personal observation and SWAO, 2023).

**Figure 1.**
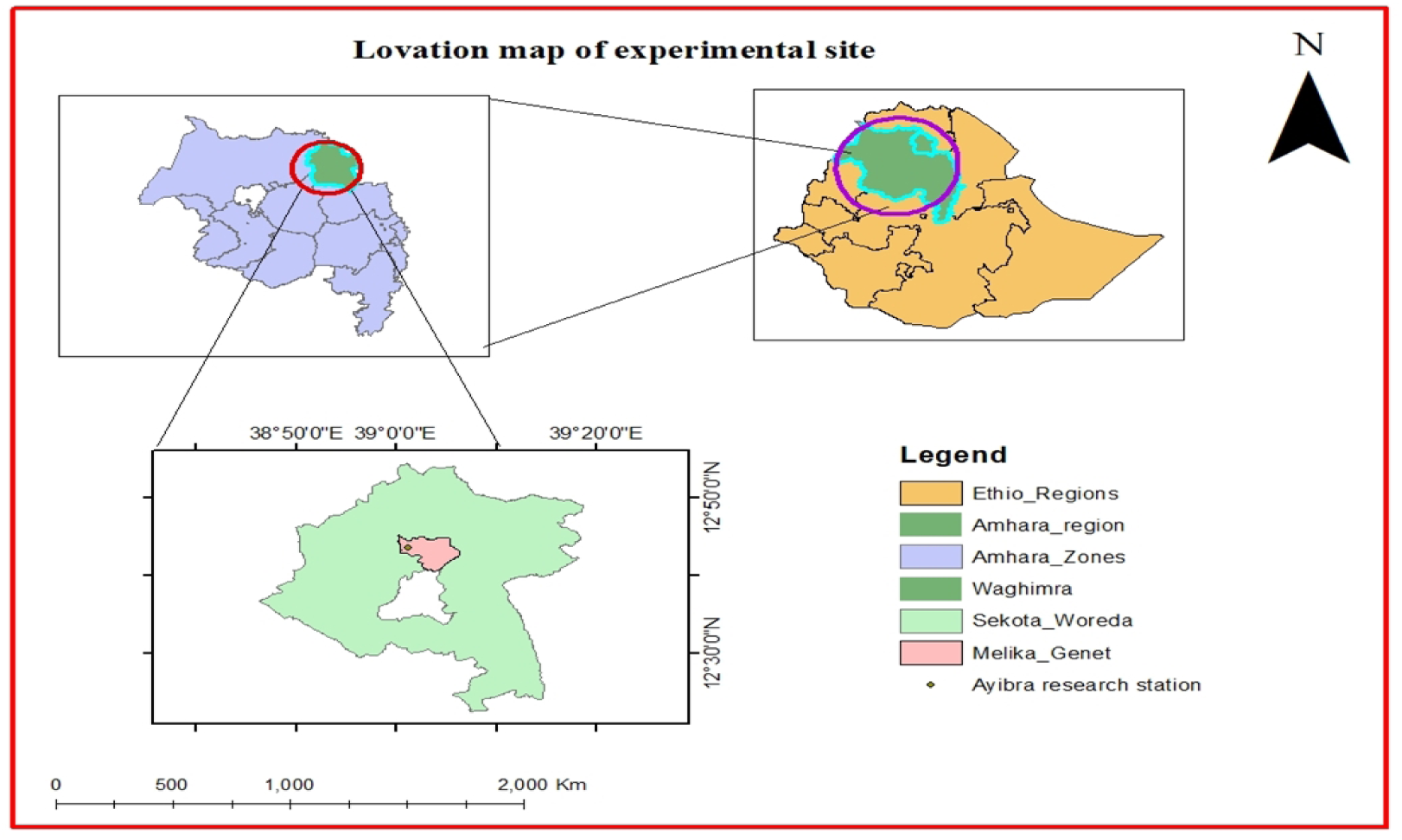
Location of the study area in the Sekota District during the 2023 cropping season.

### Experimental Materials

The experiment was conducted on sixty four small-seeded white-type common bean genotypes. Sixty-one of them are newly advanced (fixed line) genotypes that were developed by the Melkassa Agricultural Research Centre (MARC) for moisture deficit areas, and three common bean varieties namely Awash-2, Awash Melka and Awash Miten as standard checks. Both the newly developed genotypes and the checks were obtained from the National Low Land Pulse Research Coordinating Centre of the Melkassa Agricultural Research Centre (MARC). Detailed descriptions of the materials are provided in Appendix Table 1.

### Experimental Design and Management

The field experiment was conducted in a partially balanced simple lattice incomplete block design, with 8 x 8 arrangements. The individual plots size was contained rows of 2 meters (m) in length and width, each separated by 0.4 m in row spacing *i.e.* 2 m x2 m = 4 meter square (m^2^) in a grosses area and net area of 2 m x 1.2m (2.4m^2^) within a total field area of 20.2 m width x 40 m length (808 m^2^). The distances between plots, intra blocks, and replications were 0.5 m, 0.6 m, and 1 m, respectively. The treatments (genotypes) were allocated to plots at random within each intra-block and inter-block (replication).

Fertilizer was applied to all plots uniformly in the form of NPSB fertilizers at a rate of 100 kilogram per hectare and all the NPSB fertilizer were applied by side dressing during planting.

In May and late June 2023 G.C, the experimental land (plot) was prepared and uniformly labeled. Each plot was evenly seeded by hand at its blanket recommendation of spacing between plants at 0.1 cm into the slighted hole at the recommended rate of 150 kg ha^-1^. To suppress flea beetles (*Trirhabda flavolimbta)*, a two-times Karate chemical (1 L of the chemical dissolved in 200 L of water ha^-1^) was sprayed soon after the emergence of common beans when foliage beetles are seen on the leaves of the common beans. The second spray was made seven days after the first spray. All experimental plots were uniformly weeded manually at a frequency of two-time at the same time for all experimental plots.

### Data Collection and Measurement

The pre and post-harvest data on grain yield and yield-related traits of common bean genotypes were collected on a plot and plant basis following the data scoring standards of the IBPGR for common beans (IBPGR, 1985). Only three central rows were used for data collection. Five randomly selected plants from the three central rows of each plot (experimental unit) in each replicate were used for plant-based data collection after tagging. The averages of the five plants in each plot were used for statistical analysis of the traits recorded on a plant basis, while the data from all three central rows data were used for plot-based data.

#### Plot-based data

**1. Days to 50% flowering (days):** The number of days was calculated by subtracting from sowing to a stage when 50% visual judgment of the plants in a plot produced flowers.
**2. Days to 90% physiological maturity (days):** The number of days was calculated from sowing to 90% visual judgment of the plants that attained physiological maturity in each plot and when grains were difficult to divide by thumbnail.
**3. Hundred Seed weight (g):** The grain weights of 100 seeds sampled at random from the total seeds harvested from the experimental plot were counted using an electronic seed counter weighted by using an electronic sensitive balance and recorded and adjusted to a

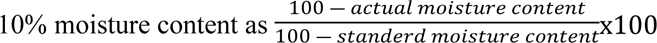

**4. Seed yield (g/plot):** The seed yield per plot after harvested, threshed, and cleaned was weight using a sensitive balance and then the moisture of the seed yield were adjusted to 10 % moisture level and converted to kg ha^-1^. Adjusted seed yield= (100-actual moisture content) / (100-standard moisture content) X obtained yield based on Hellevang (1995) formula. Yield advantage of the tested genotypes over the checks was calculated as

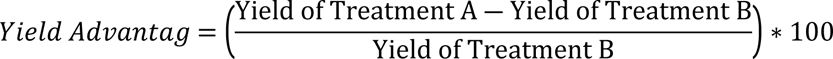

Yield of treatment A refers to the yield obtained from the treatment being compared against (yield of the new genotype) and yield of treatment B refers to the yield obtained from the treatment to which treatment A is being compared (yield of the checks used during the study).

**5. Biomass yield (g/plot):** The total biomass yield was recorded by weighing the total sun-dried above -ground biomass harvested from the three central rows and converted in to kg ha^-1^.
**6. Harvest index (%):** It was estimated by dividing the seed yield per plot by above ground biomass yield and multiplying by 100. i.e. seed yield(economic yield)/ above ground biomass x 100.

#### Plant-based data

**1. Plant height (cm):** plant height was measured in cm from ground level to the top of the plants at random from the three rows.
**2. Number of primary branch/ plant (No.):** The total number of branches per plant from the five sampled plants from the central three rows of each plot excluding the main plant, were recorded by counting at maturity
**3. Number pods per plant (No.):** In the central three rows, the total number of pods per plant at the time of harvest from five randomly selected plants is expressed as an average.
**4. Number of seeds per pod (No.):** The number of seeds per pod was recorded by counting the number of seeds per pod on each pod and was expressed as an average of five plants on a plant.

### Statistical Data Analysis

#### Analysis of Variance

The data collected for quantitative traits analysis were subjected to analysis of variance (ANOVA) for simple lattice design using the PROC LATTICE and PROC GLM procedures of SAS software version 9.2 (SAS Institute Inc., 2008) according to the procedures outlined by Gomez and Gomez (1984) after testing ANOVA assumptions and relative efficiency (RE) of design (simple lattice) over RCBD. The normality of the data was checked using the Shapiro-Wilk test before analysis. The mean squares of the traits for which RE > 100% were taken from the lattice, while those for which RE< 100% were taken from the GLM results. The general linear model (GLM) is used to calculate the unadjusted block sum of squares, unadjusted treatment sum of squares, and intra block error and handles relating one or several continuous dependent variables to one or several independent variables. Mean separations were carried out following the statistical significance of genotype mean squares using the

Duncan multiple range test (DMRT) at 5% and 1% significance levels to compare genotypes means. Estimates of genetic parameters and associations of traits (correlations) for the genotypes were continued based on the statistical significance of the mean squares of the genotypes.

The statistical model for simple lattice design:

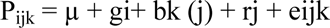

where, P_ijk_ = is the phenotypic value of the i^th^ genotype under the j^th^ replication and k^th^ incomplete block

within replication j; µ = is the grand mean constant common to all observations; gi = the effect of i^th^ genotype; bk (j) = is the effect of incomplete block k within replication j; rj = the effect of replication j; and eijk = the residual or effect of random error.

A simple lattice design is one of the twice replicated partially balanced lattice design that has four main sources of variation that can be attributed to a simple lattice design: replication, treatment (genotype), incomplete blocking, and experimental error. Relative to the RCBD design, the incomplete block is an additional source of variation and reflects the differences among incomplete blocks of the same replication.

### Estimation of Genetic Parameters

#### Estimation of phenotypic and genotypic variances

The total variance was partitioned into components due to genotype and environment. The phenotypic and genotypic variances of each trait were estimated from a simple lattice and GLM analysis of variances.

The genotypic and environmental variances for different traits were computed as per the methods suggested by Burton and Devane (1953).

Genotypic variance 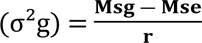 and environmental variance (σ^2^e) = MSe. Then, the phenotypic variance was obtained from the sum of genotypic and environmental variances as follows: σ^2^p = σ^2^g + σ^2^e.

Where:

σ^2^g= Genotypic variance (adjusted); σ^2^p=Phenotypic variance; σ^2^e= Environmental variance; MSe= Error mean square (Environmental variance); MSg = Mean square due to genotypes

and r = Number of replications.

Coefficients of variations at Phenotypic and genotypic levels were estimated based on the method suggested by Burton and Devane (1953) and Deshmukh *et al. (*1986) using the following formula: 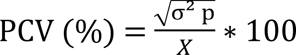 and 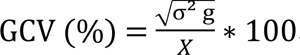. Where: PCV= Phenotypic coefficient of variation (%); GCV= Genotypic coefficient of variation (%); σ^2^p= Phenotypic variance; σ^2^g= genotypic variance and X=is the Grand mean. Genotypic coefficient of variation (GCV) and Phenotypic coefficient of variation (PCV) values were categorized as low (0-10%), moderate (10-20%), or high (20% and above) values as indicated by Burton and Devane (1953).

#### Estimate of broad-sense heritability

Heritability specifies the proportion of the total variability that is due to genetic causes. Information on heritability provides the relative practicability of selection for a particular character. Different characters have different levels of heritability that can contribute to yield improvement in breeding programs. Broad sense heritability values on plot and plant basis was computed for all traits based on the formula adopted from Johnson H*.et al*. (1955), Allard, R. (1960), and Falconer and Mackay (1996) as follows:

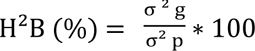

H^2^B = Heritability in a broad sense, σ^2^p = Phenotypic variance, σ^2^g = Genotypic variance and broad sense heritability estimates were classified as low (< 30%), moderate (30-60%) and high (>60%) as described by Robinson *et al*. (1949) and Falconer and Mackay (1996).

#### 3.6.3 Estimation of expected genetic advance

Although high heritability estimates have been found to be effective in the selection of superior genotypes on the basis of phenotypic performance, Johnson *et al*. (1955) suggested that heritability estimates along with genetic advances are more useful in predicting the effect of selecting the best individual. The expected genetic advance for each trait at selection intensity of 5% was computed using the methodology given by Allard (1960) and Johnson *et al*. (1955) as follows;

GA = K ∗ H^2^B ∗ σp. Where: GA= Expected genetic advance under selection, K = the standardized selection, the differential constant at 5% selection intensity (K = 2.063), σp = Phenotypic standard deviation, and H^2^B= heritability in a broad sense.

The genetic advance as a percent of the mean was calculated to compare the extent of the predicted advance of different traits under selection, using the formula given by Robinson *et al*. (1949) as follows 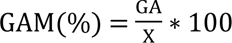. Where: GAM = Genetic advance as percentage of the mean; GA = expected genetic advance under selection and X = grand mean. The genetic advance as a percent of the mean (GAM) was categorized as low (0-10 %) moderate (10-20 %: or high (20 % and above) as given by Johnson *et al. (*1955) and Falconer and Mackay (1996).

### Estimation of Correlation (traits associations) and Path Coefficient

#### Estimation of Genotypic and Phenotypic Correlation Coefficients

Phenotypic and genotypic correlation coefficients were computed from variance and covariance components based on the method described by Singh and Chaudhury (1996). Genotypic correlations were calculated by using genotypic variances and covariance while phenotypic correlations were calculated by using phenotypic variances and covariance.

The PROC CANDISC procedure in SAS 9.2 software was used for the genotypic and phenotypic correlation significance tests. The levels of significance of the correlation coefficients were determined from the correlation table at the appropriate degrees of freedom and probability level (Gomez and Gomez, 1984). Phenotypic and genotypic correlation coefficients were computed from variance and covariance components based on the method described by Singh and Chaundry (1985) as follows.

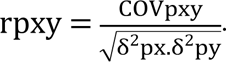

Where:

rpxy = Phenotypic correlation coefficient between traits x and y, Covpxy = Phenotypic covariance between traits x and y, δ^2^px = Phenotypic variance of trait x, δ^2^py = Phenotypic variance of trait y.

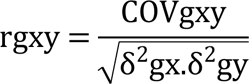

Where:, rgxy= Genetic correlation coefficient between traits x and y

Covgxy = Genetic covariance for traits x and y, δ^2^gx = Genetic variance for trait x, δ^2^gy = Genotypic variance for trait y.

The significance of the computed phenotypic correlation values was assessed using the t-test.

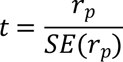

where rp =is the phenotypic correlation and SE (rp) = is the standard error of the phenotypic correlation, which were obtained via the following procedure(Sharma, 2006).

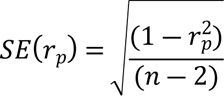

Where rp is the phenotypic correlation coefficient and n is the number of genotypes that were tested. The significance of the coefficients of correlation at genotypic levels was assessed using the formula provided in(Robertson, 1959) as shown below.

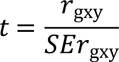

#### Path coefficient analysis

The correlation coefficient shows the relationship between variables, without identifying their direct or indirect effects. It represents the overall effect of variables on each other. Path analysis, as described by Singh and Chaudhary (1977), breaks down the overall correlation into direct and indirect effects of independent variables on the dependent variable. Since the correlation coefficient is not sufficient to explain the true relationships among traits, the path coefficient shall be worked out. In this path coefficient analysis, seed yield and seed yield-related traits were denoted as dependent (response) and independent variables, respectively (Singh and Chudhary, 1977). The direct and indirect effects of seed-related traits on seed yield were analyzed through path coefficient analysis. The analysis was computed as suggested by Dewey and Lu (1959) using the following formula

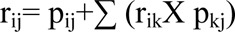

Where: r_ij_= is the Mutual association between independent traits (i) and dependent traits (j) as measured by the correlation coefficient, P_ij_= Component of the direct effect of an independent trait (i) on a dependent variable (j).

∑ r_ik_ *p_kj_ = Summation of components of the indirect effect of a given independent traits via all other independent traits. The residual effect, which determines how best the causal factors account for the variability of the dependent factor grain yield, was calculated using the formula (Dewey and Lu, 1959):

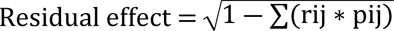

Where: p*_ij_*=Component of direct effects of the independent character (i) and dependent character (j) as it was measured by the path coefficient;

r*_ij_* =Mutual association between the an independent character (i) and a dependent character (j) as it was measured by the correlation coefficient.

### Cluster Analysis and Genetic Divergence

#### Cluster analysis

Cluster analysis is a multivariate statistical analysis technique involving partitioning a set of objects/individuals into groups so that objects within a group are more similar and objects in different groups are more dissimilar (Mahalanobis, 1936). The appropriate number of clusters was determined based on Pseudo-F and Pseudo t^2^ values using the R software core team 4.3.1 (2022). The points where local peaks of the pseudo-F value join with a small value of the pseudo-t^2^ statistic followed by a larger pseudo-t^2^ value for the next cluster combination. The mean values of data were pre standardized to mean zero and variance of one to avoid bias due to the differences in measurement scales before computing because various traits were measured by different scales.

Cluster means were calculated for individual traits on the basis of the mean performance of the genotypes included within the cluster. The genetic separation between groups was also investigated using Mahalanobis D2 statistics (1936). A dendrogram was build based on Ward’s agglomerative hierarchical classification technique with Euclidian distance as a measure of dissimilarity by using R software (R core team, 2022).

#### Genetic divergence analysis

Genetic diversity is useful tool for selecting appropriate genotypes/lines for hybridization because genetically diverged parents are expected to produce maximum heterosis and wide variability in genetic architecture (Falconer, 1960, Azad *et al*., 2012). Genetic divergence analysis quantifies the genetic distance among the selected genotypes and reflects the relative contribution of specific traits to the total divergence. D-square statistics (D^2^) developed by Mahalanobis (1936) is used to classify divergent genotypes into different groups. The extent of diversity present between genotypes was determined the extent of improvement gained through selection and hybridization. The more divergent the genotypes are, the greater the probability of improvement through selection and hybridization.

The generalized genetic distance (D^2^) between inter-clusters was calculated by the R core team procedure using the generalized Mahalanobis D^2^ statistics equation (Mahalanobis, 1936) as

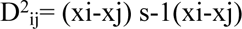

Where: D^2^ =Generalize square distance between cluster i and j xi-xj= Difference between mean vector values for i^th^ and j^th^ cluster s-1= Inverse of pooled variance covariance matrix within groups.

Average intra-cluster D^2^ values were estimated using the formula:

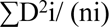

Where: ∑D^2^i = Sum of distances between all possible combinations (n) of common bean genotypes included and n=is the number of genotypes within cluster i.

The chi-squared test was used to determine the significance of the distance values. The D^2^ values obtained for all pairs of clusters are considered as the calculated values of Chi squared (Χ^2^) and were tested for significance at both the 1% and 5% probability levels against the tabulated value of Χ^2^ for the ‘P-1’ degree of freedom (DF), where P is the number of traits used for clustering genotypes (Singh and Chaudhary, 1985).

### Principal Component Analysis

Principal component analysis reflects the importance of the largest contributor to the total variation at each axis for differentiation (Sharma, 1998). A data matrix of 10 (number of traits studied) x 64 (number of genotypes used in the study) was prepared for principal component analysis. The mean values of the data were pre standardized to a mean of zero and variance of one to avoid bias due to the differences in measurement scales before computing the principal component analysis because various traits were measured at different scales. Principal component analysis was performed using R Software (R core team, 2022) to determine the contribution of each quantitative trait to the total variation in the genotypes. Therefore, the principal components (PCs), with eigenvalues greater than 1 explained by the total variation among genotypes for all traits were retained. The traits that contributed to the most to the total variation were identified in each principal component.

The principal component variables were defined as linear combinations of the original variables X _1_… X_k…_ X_m_. The extracted eigenvector table provides coefficients for equations below (Holland, 2008) as Y_k_ = C_k1_ X_1_ + C_k2_ X_2_ + …. + C_km_ X_m_, where Y_k_ is the k^-th^ principal component k, and C‘s are the coefficients in the table.

Standard criteria that permit us to ignore components whose variance explained is less than 1 when a correlation matrix is used for determining the number of PCs were employed (Holland, 2008). As suggested by Johnson and Wichern (1988), an eigenvector greater than half divided by the standard deviation (square root) of the eigenvalues of the respective PC was employed as a general guideline for weighing the relative significance of traits constituting the PCs. Principal components (PCs) with eigenvalues greater than unity, and component loadgings greater than ± 0.3 were considered to be meaningful and valuable (Hair *et al.,* 1998; Daniel *et al.,* 2011; Adebisi *et al.,* 2013).

## RESULTS AND DISCUSSION

Results obtained on variability assessment and associations among yield and yield related characters and genetic divergence are presented here under. Implications of such studies in common bean improvement and breeding program for higher seed yield and other traits of interest are also discussed.

### Analysis of Variance

The results of the analysis of variance (ANOVA) for 10 traits are presented in Table 1. There were highly significant (P˂0.01) differences among the tested common bean genotypes for all traits in this study. This justified the need to estimate statistical analyses such as genetic parameters, divergence, and correlation coefficients for the genotypes. The findings of highly significant differences among the genotypes for traits under study suggest the existence of sufficient genetic variability in the materials used for this study. In line with the present findings, several researchers have reported the existence of significant differences in traits among common bean genotypes studied for the traits (Kedir Shafi *et al.,* 2019; Nigussie Kefelegn *et al*., 2020; Belay Asmare *et al.,* 2023; Kedir Yimam *et al.,* 2023) on common bean genotypes for days to 50% flowering, days to 90% physiologically maturity, for the number of branches per plant, above ground biomass, seed yield and harvest indexes in different parts of Ethiopia.

**Table 1.**
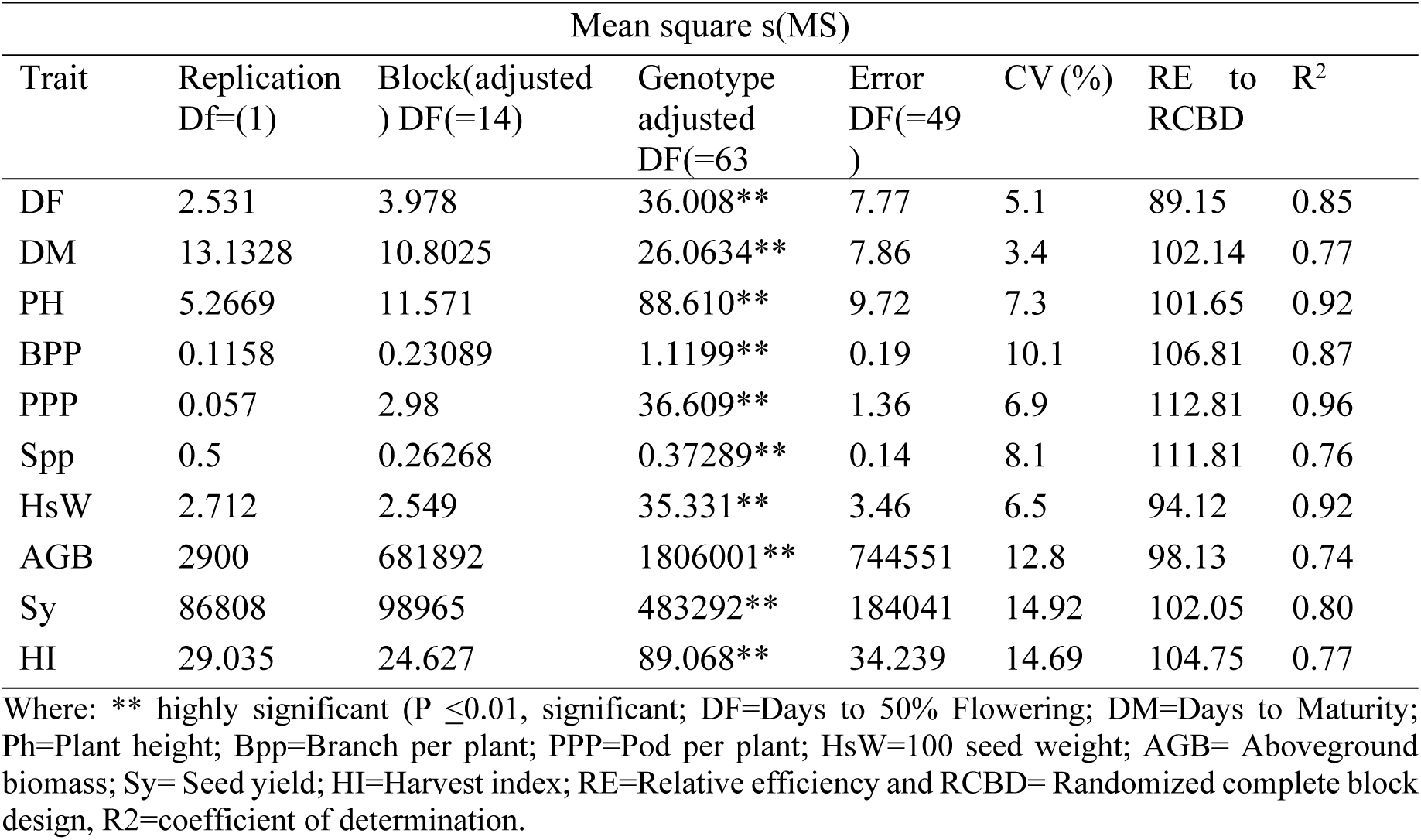
Mean squares (MS) from Analysis of variance (ANOVA) for 10 quantitative traits on 64 common bean genotypes.

| Trait | Mean square s(MS) |  |  |  |  |  | R<sup>2</sup> |
| --- | --- | --- | --- | --- | --- | --- | --- |
|  | Replication<br>Df=(1) | Block(adjusted<br>) DF=(14) | Genotype<br>adjusted<br>DF=(63) | Error<br>DF=(49) | CV (%) | RE to<br>RCBD |  |
| DF | 2.531 | 3.978 | 36.008** | 7.77 | 5.1 | 89.15 | 0.85 |
| DM | 13.1328 | 10.8025 | 26.0634** | 7.86 | 3.4 | 102.14 | 0.77 |
| PH | 5.2669 | 11.571 | 88.610** | 9.72 | 7.3 | 101.65 | 0.92 |
| BPP | 0.1158 | 0.23089 | 1.1199** | 0.19 | 10.1 | 106.81 | 0.87 |
| PPP | 0.057 | 2.98 | 36.609** | 1.36 | 6.9 | 112.81 | 0.96 |
| Spp | 0.5 | 0.26268 | 0.37289** | 0.14 | 8.1 | 111.81 | 0.76 |
| HsW | 2.712 | 2.549 | 35.331** | 3.46 | 6.5 | 94.12 | 0.92 |
| AGB | 2900 | 681892 | 1806001** | 744551 | 12.8 | 98.13 | 0.74 |
| Sy | 86808 | 98965 | 483292** | 184041 | 14.92 | 102.05 | 0.80 |
| HI | 29.035 | 24.627 | 89.068** | 34.239 | 14.69 | 104.75 | 0.77 |

Where: \*\* highly significant ( $P \leq 0.01$ , significant; DF=Days to 50% Flowering; DM=Days to Maturity; Ph=Plant height; Bpp=Branch per plant; PPP=Pod per plant; HsW=100 seed weight; AGB= Aboveground biomass; Sy= Seed yield; HI=Harvest index; RE=Relative efficiency and RCBD= Randomized complete block design, R2=coefficient of determination.

### Mean Performance for Measured Agronomy Traits

The mean values of 64 small-seeded common genotypes for each trait are presented in Appendix Table 2. The genotypes took 44-62 days on average to reach 50% flowering. Genotype number 61 took the longest (62) days to flower while genotype 29 took the shortest (44) days to flower with a mean of 54.4 ± 1.87 days. The longest and shortest 90% days to maturity were recorded for genotype number 10 (95.0 days) and for genotype number 27 (72.5 days), respectively, with an overall average of 82.2 ± 2.0 (Appendix Table 2) indicating that the tested genotypes belonged to the early and middle maturation categories. The seed yield of 64 small-seeded common bean genotypes was in the ranged from 1633.1 kg ha^-1^ to 3702.10 kg ha^-1^ with a mean of 2542.53 kg ha^-1^. In this study, the highest seed yield per hectare was recorded for genotype number 9 (3702.10 kg ha^-1^), while the lowest yield was obtained for genotype number 63 (1633.1 kg ha-1) (Appendix Table 2). The percentage of the genotypes with above average mean (2542.53 kg ha^-1^) seed yield was 45.31% (29 genotypes). Among the 61 newly fixed-line small-seeded common bean genotypes tested, 18 genotypes (29.5%) namely genotype number 2, 6, 9, 13, 14, 18, 22, 23, 24, 27, 30, 31, 34, 48, 50, 53, 56 and 60 had greater yields than did the best yielder standard check (genotype number 62, 2792.7 kg ha-1) and other tested genotype.

The yield of these genotypes was up to 24.56% greater than that of the best seed yielder standard check (genotype number 62, Awash-2) 38.90% over the medium seed yielder standard check (genotype number 64, Awash Melka) and 55.89% over the lowest seed yielder standard check (genotype number 63, Awash Miten). Out of top yielder genotypes, five genotypes (8.20%) are early maturing namely genotype number. This suggests the chance of selecting both high yielder and early maturing genotypes that can escape terminal moisture stress. Similarly, in a study by Kedir Yimam *et al. (*2023) on 64 small-seeded common bean genotypes at Melkassa, found significant variation in seed yield ranging from 1685.09 to 4499 kg ha^-1^ with a mean of 2911.04 kg ha^-1^. In agreement with these findings, significant yield variation among 64 small-seeded common bean genotypes ranging from 980.5-2242.0 kg ha^-1^ within an average of 1675 kg ha^-1^ at Goro and 1479 kg ha^-1^ at Ginnir (Bale and East Bale Zones) was reported by Belay Asmare *et al*. (2023), and also in agreement with the present findings, Girum Kifle (2019) reported a significant difference in seed yield ranging from 1668.8 kg ha^-1^ to 4014.7 kg ha^-1^ with a mean of 2863.7 kg ha^-1^ in thirty common bean genotypes studied in the different parts of Ethiopia (Melkassa, Alemtena, Arsinegele, Haramaya University and Sirinka). In line with this, significant seed yield variation ranged from 1052 to 4456 kg ha^-1^ with an average yield of 3318 kg ha^-1^ from 64 common bean genotype studies at Haramaya University (Dire Dawa) as reported by (Yonas Moges, 2017). Kedir Yimam *et al*. (2023) reported a variation in the number of seeds per pod ranging from 4.13 to 6.66 in common bean genotypes and Chaudhary *et al*. (2020) reported a variation in the number of seeds per pod ranging from 2.07-14.43 in the number of seeds per pod in common bean genotypes. Overall, the range (the highest and lowest values) and mean value indicated that there was a considerable amount of diversity among the common bean genotypes under investigation for the majority of the attributes. As recommendation, these high-yielding and early maturing genotypes could be selected for further breeding programs (further testing over years and across locations) for the study area under rain-fed conditions. Genotypes number 27 and 30 were both early flowering, early matured and higher in seed yield than the average mean and the checks. Therefore using those genotype (genotype number 27 and 30) as parents in crosses will help to improve days to flowering, days to maturity and seed yield at the same time.

### Estimates of Variance Components

#### Phenotypic and genotypic variations

The estimated variance components, PCV, GCV, HB2 and GAM values of the traits considered in this study are presented in Table 2.

**Table 2.** Minimum, maximum, mean, variances, coefficients of variation, heritability and genetic advance as percentages of the means of 10 traits of small-seeded common bean genotypes.

| Trait | Minimum | Maximum | Mean | $\sigma^2_g$ | $\sigma^2_p$ | $\sigma^2_e$ | GCV<br>(%) | PCV<br>(%) | H2B<br>(%) | GA | GAM<br>(5 %) |
| --- | --- | --- | --- | --- | --- | --- | --- | --- | --- | --- | --- |
| DF | 43.00 | 64.00 | 54.4375 ± 1.87 | 14.54 | 21.47 | 6.93 | 7.00 | 8.51 | 67.7 | 6.46 | 11.88 |
| DM | 71.00 | 96.00 | 82.2422 ± 2.0 | 8.77 | 17.29 | 8.51 | 3.60 | 5.06 | 50.75 | 4.35 | 5.29 |
| PH | 26.23 | 61.80 | 42.8018 ± 2.23 | 39.24 | 49.37 | 10.13 | 14.63 | 16.42 | 79.47 | 11.50 | 26.88 |
| BPP | 2.20 | 5.80 | 4.3043 ± 0.32 | 0.46 | 0.66 | 0.20 | 15.76 | 18.87 | 69.8 | 1.17 | 27.13 |
| PPP | 9.40 | 30.80 | 16.8633 ± 0.93 | 17.44 | 19.16 | 1.72 | 24.77 | 25.96 | 91.02 | 8.21 | 48.68 |
| Spp | 3.00 | 5.40 | 4.3578 ± 0.28 | 0.11 | 0.26 | 0.15 | 7.58 | 11.78 | 41.37 | 0.43 | 10.04 |
| HsW | 18.00 | 40.20 | 28.6677 ± 1.28 | 16.03 | 19.30 | 3.26 | 13.97 | 15.32 | 83.09 | 7.52 | 26.23 |
| AGB | 4500.00 | 9804.35 | 6730.5929 ± 604.4 | 53768313.6 | 1268313 | 730626.5 | 10.89 | 16.73 | 42.39 | 983.5 | 14.61 |
| Sy | 1484.23 | 3974.11 | 2542.5345 ± 266.3 | 170744.92 | 312547.24 | 141802.3 | 16.25 | 21.99 | 54.63 | 629.2 | 24.75 |
| HI | 21.00 | 58.96 | 38.2064 ± 4.0 | 28.48 | 60.59 | 32.10 | 13.97 | 20.37 | 47.01 | 7.54 | 19.73 |

Where $\sigma^2_g$ =Genotypic variance; $\sigma^2_p$ = Phenotypic variances; GCV (%)= Genotypic coefficient of variation; PCV(%)=Phenotypic coefficient of variation; H2B (%)= Broad sense heritability; GA=Genetic advance; GAM (%)=Genetic advance as percent of mean; Df= Days to 50% flowering; DM= Days to 90% maturity; Ph=Plant height (cm); BPP branch per plant (no); PPP= pod per plant (no); SPP=seed per pod (no); HsW=100 seed- weight (gr); AGB= Above ground biomass (kg ha<sup>-1</sup>); Sy= Seed yield (kg ha<sup>-1</sup>) <sup>1)</sup> and HI=Harvest index (%)

The greatest value of phenotypic variance was calculated for plant height (49.37) while the lowest was for the number of seeds per pod (0.26). Genotypic variances ranged from 0.11 for seeds per pod to 53768313.55 for aboveground biomass. The environmental variances were also relatively greater for traits above ground biomass (730626.50), seed yield (141802.32), harvest index (32.10), and plant height (10.13) than for the other traits. This indicates that there is a considerable influence of environmental factors on the phenotypic expression of these traits. This result is in agreement with the findings of Ejigu Ejara *et al*. (2018), Kedir Shafi *et al*. (2019), Masreshaw Yirga *et al*. (2022), and Kedir Yimam *et al*. (2023), who reported relatively high phenotypic variance for days to 50% flowering, days to 90% maturity, seed per pod, biological yield, harvest index and seed yield and other studied traits in different parts of Ethiopia for different numbers of common bean and soybean genotypes.

The estimated genotypic coefficient of variation (GCV) values ranged from 3.6% for days to 90% maturity to 24.77% for pods per plant while the phenotypic coefficient of variation (PCV) ranged from 5.06% for days to 90% maturity to 25.96% for branches per plant (Table 2). The number of pods per plant, (25.96), seed yield (21.99), and harvest index (20.37) had high PCV while the number of branches per plant (18.87), plant height (16.42), and number of seeds per pod (11.78), hundred seed weight (15.32), and aboveground biomass (16.73) had moderate PCV. However, days to 50% flowering (8.51) and 90% maturity (5.06) had the lowest PCV, suggesting the presence of sufficient variation among the tested genotypes and indicating a better response to selection.

In the present study, pods per plant (24.77) had the highest GCV, while Plant height (14.63), branches per plant (15.76), hundred seed weight (13.97), aboveground biomass (10.89), seed yield (16.25) and harvest index (13.97) had moderate GCV. The number of days to 50% flowering (7.00), days to 90% maturity (3.60), and number of seeds per pod (7.58) had lower GCV. Traits that have low GCV indicate less scope of selection due to the absence of sufficient variation among tested genotypes and indicate the need for the creation of variability through different breeding methods such as introduction, collection, hybridization, mutation *etc.* Low to high GCV and PCV indicate that there were considerable variations among environmental factors between replications and genotypes. In line with these findings, Similarly, Nigussie Kefelegn, *et al*. (2020), on 15 common bean genotypes at Chefa, report a high value of PCV for pod per plant, seed yield, and harvest index. Also in line with these findings, Meseret Tola, *et al*. (2020) on 25 common bean genotypes at Melkassa, and Yonas Moges (2017) on 64 common bean genotypes at Dire Dawa reported moderate PCV for hundred seed weight, seed per pod, and aboveground biomass. In contrast to these findings, high PCV value were estimated for aboveground biomass, plant height, number of seeds per pod, number of branches per plant, and hundred-seed weight for 100 soybean genotypes at Pawe was reported by (Masreshaw Yirga *et al*., 2022), and low PCV were estimated for seed yield, hundred-seed weight, and number of pods per plant by Henry *et al*. (2019) for 52 common bean genotypes in Kenya. In line with this result, low PCV for days to flowering and days to maturity was reported by Nigussie Kefelegn *et al*. (2020) and Yonas Moges (2017). Moderate GCV for hundred seed weight, harvest index, and plant height were reported by Tekalign Afeta *et al*. (2020) and Desalegn Nigatu *et al*. (2022). Similarly, Simon Yohannes *et al*. (2020) reported high GCV for pods per plant on 33 common bean genotypes in Areka, southern Ethiopia, and in contrast to these findings by the same authors, high GCV for seed yield, hundred seed weight, moderate GCV for seeds per pod and low GCV for harvest index were also reported. Additionally, Nigussie Kefelegn *et al*. (2020) reported a moderate GCV for plant height on 15 common bean genotypes in Koga by. Consistent with these findings, Belay Asmare *et al*. (2023) reported moderate GCV on plant height, hundred-seed weight, and seed yield in 64 small-seeded common bean genotypes in Goro and Ginnir, Southeast Ethiopia. Moderate GCV for seed yield was also observed by Sima Jibril *et al*. (2019) for 20 common bean genotypes in Haramaya, Eastern Ethiopia.

Similarly, low GCV for days to flowering and days to maturity have been reported by different authors, such as Tekalign Afeta *et al*. (2020), Owusu *et al*. (2021), Desalegn Nigatu, *et al*. (2022), Belay Asmare *et al*. (2023) and Kouam *et al*. (2023). In contrast, high GCV for days to flowering was reported by Mofokeng *et al. (*2020), and moderate GCV for days to maturity was reported by Henry *et al*. (2019).

The differences in magnitude between PCV and GCV for the following traits were relatively high: number of branches per plant, number of seeds per pod, aboveground biomass, seed yield, and harvest index were relatively high compared to other traits. A greater PCV than GCV for the studied traits implies that the influence of environmental factors on the phenotypic expression of these traits and the variation among the genotypes is not solely due to genotypic expression. This may make it difficult or practically impossible to improve the characters by direct selection of high-performing genotypes. However, the differences between the PCV and GCV values were small for days to 50% flowering, 90% days to maturity, the number of pods per plant, plant height, and hundred-seed weight. This minimum difference between PCV and GCV tells as that more of the phenotypic expression came from genotypic influences and that the environmental influence was also less. Their phenotypic expression would be a good indication of genetic potential and the effectiveness of the selection of genotypes on the basis of the phenotypic performance of these traits. Thus, these characteristics could be used as direct selection criteria for common bean improvement in the studied area. Similarly, a greater PCV than GCV in trait studies was reported by Meseret Tola *et al*., 2020; Tekalign Afeta *et al.,* 2020; Owusu *et al.,* 2021; Tekle Yoseph *et al*., 2022; Lenjisa Tamiru *et al*., 2023).

#### Heritability and genetic advance

The estimated broad-sense heritability values for the traits studied are presented in Table 2. Medium to high heritability values were observed for the traits under study and it ranged from 0.41 (seed per pod) to 0.91 (pod per plant). High estimates of broad sense heritability were observed for days to 50% flowering (68%), plant height (79%), number of branches per plant (70%), hundred seed weight (83%), and number of pods per plant (91%). Moderate heritability values were recorded for days to 90% maturity (51%), number of seeds per pod (41%), aboveground biomass (42%), seed yield (55%), and harvest index (47%).and these high heritability values indicate a high level of genetic variance as a component of the total variance or variation observed, mainly under genetic control. In other words, these traits are less influenced by environmental factors in their expression and heritable/inherent potential differences among the traits. High heritability values for days to flowering, and hundred-seed weight, and moderate heritability values for the number of seeds per pod and seed yield have been reported previously with the heritability for days to maturity contradicts the previous findings by Eyuel Mesera *et al*. (2022). In line with this, high the broad-sense heritability of plant height and hundred seed weight was reported before Kedir Gelgelu *et al.,* 2017; Awol Mohammed *et al.,* 2019; Ghimire and Mandal 2019; Fantaye Belay and Kibrom Fisseha 2021). In contrast with this low heritability value in broad sense for hundred seed weight were done by Mesfin Hailemariam. (2018).

Similar, moderate heritability values for the number seeds per pod, aboveground biomass, and harvest index were also reported by Kedir Gelgelu *et al*. (2017), Meseret Tola *et al*. (2020), and Belay Asmare *et al*. (2023). In addition, in line with these findings, moderate heritability in seed yield was reported by (Mofokeng *et al.,* 2020; Desalegn Nigatu *et al*., 2022; Masreshaw Yirga *et al*., 2022 and Tekle Yoseph *et al*., 2022). However, in contrast to this result, high heritability value in the broad sense for seed yield was recorded by (Chaudhary *et al*., 2020; Nigussie Kefelegn *et al.,* 2020; Owusu *et al.,* 2021; Kouam *et al*., 2023). Similar to these findings, a moderate heritability value for days to 90% maturity was reported by Tekalign Afeta *et al.,* (2020). In general, moderate to high heritability in a broad sense among the studied traits indicated that the heritable nature of the traits differs and that the influence of the environment on each trait was also varied.

The estimated genetic advance as a percentage of the mean for the traits considered in the 64 small-seeded common bean genotypes studied is presented in Table 2. Genetic advance as a percentage of the mean (GAM) at 5% selection intensity (k=2.063) ranged from 5.29% for days to 50% maturity to 48.68% for the number of pods per plant, indicating that selection of the top 5% of the base population could result in an advance of 5.29 (days to 90% maturity) to 48.68 (pods per plant) percent over the respective population mean. In this study, GAM (5%) was high for seed yield (24.75%), hundred seed weight (26.23%), number of pods per plant (48.68%), number of branches per plant (27.13), and plant height (26.88%). A moderate GAM (5%) was computed for days to 50% flowering (11.88%), aboveground biomass (14.61%), and harvest index (19.73%). A low GAM (5%) was observed for days to 90% maturity (5.29%), and the number of seeds per pod (10.04 %). Moderate to high GAM indicating that these traits are governed by additive gene action and a fixable nature; hence, direct improvement of the traits through direct selection for these traits is effective. Similarly, high GAM for seed yield, hundred-seed weight, plant height, and number of pods per plant were reported by (Chaudhary *et al*., 2020; Mofokeng *et al.,* 2020; Nigussie Kefelegn *et al.,* 2020; Esrael Kanbata, 2023 and Kedir Yimam *et al.,* 2023). Low GAM for days to maturity was also reported by Mesfin Hailemariam (2018), Awol Mohammed *et al*. (2019), Amare Tsehaye *et al*. (2020) and Tekalign Afeta *et al*. (2020). In Contrast a high GAM for the number of seeds per pod was reported by Chaudhary *et al*. (2020), and a moderate GAM for number of seeds per pod was reported by Owusu *et al*. (2021).

In the present study, high heritability associated with high GAM was estimated for the plant height (0.79, 26.88), number of branches per plant (0.7, 27.13), pod per plant (0.91, 48.68), and hundred seed weight (0.83, 26.23) respectively (Table 2), suggesting the importance of additive gene action in the expression of these traits. This provides a chance for further improvement in advanced generations through selection. High estimates of GAM coupled with moderate heritability were observed for seed yield (24.75, 55%), and high heritability coupled with moderate GAM was estimated for days to 50% flowering (68, 11.88%), while moderate heritability with moderate genetic advance was noted for the aboveground biomass (0.42, 14.61%) and the harvest index (0.47, 19.73). High heritability coupled with high GAM for the traits of the number of pods per plant, hundred seed weight, and number of branches per plant was also reported by Awol Mohammed *et al*. (2019), Kedir Shafi *et al*. (2019), Henry *et al*. (2019), Tekalign Afeta *et al*. (2020) and Belay Asmare *et al*. (2023). High heritability in the broad sense with a high GAM for hundred seed weight and plant height was also reported by (Sima Jibril *et al*. (2019), Owusu *et al*. (2021), and Fantaye Belay and Kibrom Fisseha (2021).

Similarly, high GAM and moderate heritability in the broad sense of seed yield were reported by (Sima Jibril *et al.,* 2019; Desalegn Nigatu *et al*., 2022; Masreshaw Yirga *et al*., 2022; Tekle Yoseph *et al*., 2022; Kedir Yimam *et al.,* 2023). On the other hand, moderate heritability with low genetic advancement as a percentage of the mean was exhibited for days to 90% maturity (51%, 5.29%), and for seeds per pod (41%, 10.04%). The lower values of heritability and GAM may be due to non-additive gene action and greater environmental factors according to Falconer and Mackay (1996). This suggests that non-additive gene action predominates in the expression of these traits, which can be exploited by heterosis breeding and recurrent selection (Panse, 1957). Furthermore, the most reliable estimate of the degree of genetic advance through phenotypic selection is provided by high heritability combined with high estimates of the genetic coefficient of variation (Burton, 1952). Therefore, selection may be effective in early generations in which the selection of genotypes can be based on phenotypic performance (Johnson *et al.,* 1955). Based on this suggestion, a high genotypic coefficient of variation (GCV) with a high heritability value and a high GAM were estimated for the trait pod per plant (24.77, 91 and 48.68%) respectively. Similarly, high GCV, heritability and GAM for pods per plant were reported by (Amare Tsehaye *et al.,* 2020; Simon Yohannes *et al.,* 2020; Masreshaw Yirga *et al.,* 2022; and Welde Ketema and Negash Geleta 2022). In contrast with this finding, moderate GCV, and heritability coupled with moderate GAM for pods per plant were reporetd by (Meseret Tola *et al*. (2020). Moderate GCV was associated with high H2B and high GAM for plant height, number of branches per plant, and hundred-seed weight (Table 2). In line with these findings, moderate GCV, high heritability, and high GAM for plant height and hundred seed weight were reported by Belay Asmare *et al. (*2023) and Kedir Yimam *et al*. (2023). Moderate GCV, high heritability, and high GAM indicate that the variation is due to genetics, a highly heritable nature, and controlled by additive gene action. These traits could be considered the best components for direct selection breeding for common bean improvement. Therefore, direct selection of the best genotypes based on these traits is possible in the early generations and rewarding. In this study, moderate GCV, moderate and moderate H2B with high GAM were used for seed yield, and moderate GCV, and moderate H2B, and moderate GAM were for the harvest index. In line with these findings, moderate GCV, and moderate H2B with high GAM for seed yield were estimated by Masreshaw Yirga *et al*. (2022) and Kedir Yimam *et al. (*2023).

### Phenotypic and Genotypic Correlation

#### Phenotypic correlations of seed yield with other traits

The phenotypic correlation coefficients between seed yield and seed yield-related traits ranged from −0.09 to 0.69 (days to flowering to harvest index). Seed yield (kg ha^-1^) had a significant (p < 0.05) positive phenotypic correlation with the number of branches per plant (r_p_= 0.197*) and highly significance (p < 0.01) correlation with aboveground biomass (r_p_ = 0.44**), and harvest index (r_p_= 0.69**) (Table 3). This indicates that genotypes with a maximum branch per plant, high biological yield (biomass), and high harvest index had greater seed yields than those with a lower number of branches per plant, low biological yield, and harvest index. This may be that physiologically high biomass may correlated with seed yield due to high in branches, or pod formation. Therefore, any improvement in these traits would lead to a substantial increase in common bean seed yield. In line with these findings, (Awol Mohammed and Asnake Fikre 2018; Desalegn Nigatu *et al*., 2022; Eyuel Mesera *et al*., 2022; Esrael Kanbata 2023; and Kedir Yimam *et al*., 2023) reported a significant and positive relationship between seed yield and the biomass yield and harvest index at the phenotypic levels for cowpea and common bean genotypes. Similarly, significant positive correlations between the number of branches per plant and seed yield at the phenotypic level were reported by Itefa Degefa *et al*. (2014), Ejigu Ejara *et al*. (2017), Kumar (2023), and Belay Asmare *et al*. (2023). A positive and non-significant association of seed yield was observed with plant height (r_p_= 0.17), plant per pod (r_p_= 0.08), and seed per plant (r_p_= 0.07), while a negative non-significant phenotypic correlation of seed yield and hundred seed weight (r_p_= −0.06) days to 50% flowering (rp= −0.09) and 90% days to maturity (rp= −0.05) was observed. Similarly to this, a negative phenotypic correlation between seed yield and hundred seed weight was reported by Lenjisa Tamiru *et al*. (2023) and a negative correlation between seed yield with days to maturity was observed by Girum Kifle *et al*. (2019), Kedir Shafi *et al*. (2019), and Chaudhary *et al*. (2020). Additionally, Wakjira Getachew *et al*. (2021) reported a negative correlation between seed yield and days to flowering. Abel Moges *et al*. (2017) confirmed a negative phenotypic correlation of seed yield with days to flowering, days to maturity and 100-seed weight on 25 common bean genotypes for 11 qualitative and quantitative traits in different parts of Ethiopia (Melkassa, Srinka, Hawassa, Jimma, Arsinegelle, Areka and Alemtena) implying that an increase in days to flowering and days to maturity, may decrease seed yield. But contrasting with this, a positive significant correlation of seed yield with days to maturity was reported by Ghimire and Mandal (2019). The presence of a highly significant and positive correlation of these traits with seed yield at the phenotypic level indicated the importance of these traits in selection programs for identify common bean genotypes with high seed yield (Falconer and Mackay 1996).

**Table 3.** Estimates of phenotypic (below diagonal) and genotypic (above diagonal) correlation coefficients among 10 traits in 64 small seeded common bean genotypes.

| Traits | DsF | DsM | Ph | BPP | PPP | Spp | HsW | AGB | Sy | HI |
| --- | --- | --- | --- | --- | --- | --- | --- | --- | --- | --- |
| DF | **1\*\*** | 0.18ns | 0.1ns | 0.30* | 0.10ns | 0.15ns | 0.03ns | 0.018ns | 0.03ns | .004ns |
| DM | 0.097ns | **1\*\*** | 0.5** | 0.34** | 0.09ns | 0.27* | 0.13ns | 0.02ns | 0.30* | 0.29* |
| PH | 0.11ns | 0.23* | **1\*\*** | 0.13ns | 0.12ns | 0.08ns | 0.21ns | 0.11ns | 0.06ns | 0.09ns |
| BPP | 0.27** | 0.2* | 0.17ns | **1\*\*** | 0.029* | 0.26* | 0.33** | 0.27* | 0.19ns | 0.013ns |
| PPP | 0.12ns | 0.12ns | 0.12ns | 0.23** | **1\*\*** | 0.05ns | 0.30* | 0.05ns | 0.078ns | 0.13ns |
| Spp | 0.16ns | 0.08ns | 0.87ns | 0.28** | 0.06ns | **1\*\*** | 0.07ns | 0.10ns | 0.02ns | 0.12ns |
| HsW | 0.06ns | 0.07ns | 0.18* | 0.29** | 0.28** | 0.029ns | **1\*\*** | 0.13ns | 0.16ns | 0.32* |
| AGB | 0.02ns | 0.007ns | 0.0022ns | 0.19* | 0.015ns | 0.056ns | 0.106ns | **1\*\*** | 0.50** | 0.19ns |
| Sy | 0.09ns | 0.05ns | 0.17ns | 0.197* | 0.08ns | 0.07ns | 0.06ns | 0.44** | **1\*\*** | 0.74** |
| HI | 0.08ns | 0.034ns | 0.15ns | 0.05ns | 0.09ns | 0.15ns | 0.17ns | 0.34** | 0.67** | **1\*\*** |

Where: \*\*, \*and ns; significant at 1% ,5% probability level and non- significant, respectively; DsH= Days to flowering (No.); DsM= Days to maturity(No.); PH=Plant height (cm); Bpp= pod per plant; Spp= seed per pod; HsW=100- seed weight; AGB= Above ground biomass (kg ha<sup>-1</sup>); Sy =seed yield (kg ha<sup>-1</sup>); HI=Harvest index (%).

#### Phenotypic correlations among other traits

The seed yield-related traits exhibited varying relations of phenotype correlation among themselves (Table 3).

Days to 50% flowering had positive non-significance correlations with days to maturity (rp= 0.097), but negative and significant correlation with branches per plant (rp= −0.27**), and negative non-significance with plant height (rp=-0.11), pods per plant (rp= −0.14ns), seeds per pod, (rp=-0.16), hundred seed weight (rp=-0.056), biological yield (rp= −0.012), seed yield (−0.09) and harvest index (rp= −0.078) (Table 3). In an argument with this, Lenjisa Tamiru *et al*. (2023) investigated a positive correlation between days to flowering and days to maturity. This indicates that late-flowering genotypes were also late-maturing, tall in plant height, late in flowering date and, late in flowering date were low in seed per pod, pod per plant as well as biological and seed yield. A negative correlation of days to flowering with branch per plant was reported by Ejigu Ejara *et al*. (2018). Similarly, Itefa Degefa *et al*. (2014) and Girum Kifle *et al*. (2019) found a negative correlation between days to flowering and hundred seed weight. But contrasting with this, positive correlations of days to flowering with 100-seed weight and seeds per pod were done by Asmamaw Amogne *et al*. (2020).

The number of days to maturity exhibited positive non-significant phenotypic correlations with pods per plant (rp=0.10) and a negative significance phenotypic correlation with plant height (rp= −0.23*) and branches per plant (rp= −0.19*) and a non-significance with other traits. This indicates that delays in maturity are associated with an increase in plant height and the number of branches per plant. In line with this, a positive correlation of days to maturity with pod per plant was done by Girum Kifle *et al*. (2019) and a negative phenotypic correlation between days to maturity with plant height was reported by Meseret Tola *et al*. (2020). However, in contrast Abel Moges *et al. (*2017) reported a significant positive phenotypic correlation between days to maturity and plant height.

Plant height had a significant positive correlation with 100-seed weight (rp= 0.18*), and there were no significant correlations between plant height and the remaining seed yield component traits (Table 3). The number of branches per plant showed highly significant positive correlations with the number of pods per plant (rp= 0.23**), number of seed per pod (rp=0.28**), and 100-seed weight (rp=0.29**), and significant positive correlations with the biological yield (rp= 0.19*), and seed yield (rp= 0.197) indicating that as the number of branch per plant increased, there were simultaneous increase in the number of pod per plant, in the number of seeds per pod, hundred-seed weight, and biological and seed yield. However, the number of branches per plant had a highly significant negative correlation with the number of days to flowering (rp= −0.27**) and negative significant correlation with days to maturity (rp= −0.2*).The number of pods per plant had a highly significant positive phenotypic correlation with the number of branches per plant (rp=0.23**) and 100-seed weight (rp=0.29**), while for the remaining traits, there was no significant correlation, indicating a simultaneous increase in number of pods per with days to flowering, maturity, seed per pod, biological, yield seed yield and harvest index impossible. In line with this, a positive phenotypic correlation between pods per plant with branch per plant was reported by Asmamaw Amogne *et al*. (2020). However, these results disagree with this, Girum Kifle *et al. (*2019) and Simon Yohannes *et al*. (2020) reported a negative phenotypic correlation btween the number of pod per plant and the hundred seed weight.

The number of seeds per pod had a highly significant positive phenotypic correlation with the number of branches per plant (rp= 0.28**) indicating a simultaneous increase in the number of branches per plant and the number of seeds per pod. However, this trait had a non-significant phenotypic correlation with the remaining studied traits. Similarly, Belay Asmare *et al*. (2023) reported a significant correlation between seed per pod with branch per plant but a non-significant between seed per pod with pod per plant, plant height, and hundred seed weight. Contrasting with this, Kedir Shafi *et al. (*2019) reported a positive significance correlation between the number of seeds per pod and days to flowering, days to maturity pod per plant and seed yield at the phenotypic level.

The hundred seed weight had a highly significant positive phenotypic correlation with branches per plant (rp=0.29**), pods per plant (0.28**), and significant correlation with plant height (0.18*) indicating a simultaneous increase in hundred seed weight with branches per plant, pods per plant and plant height together. However, this trait had non-significant phenotypic correlation with the remaining study traits. In line with these findings, Ejigu Ejara *et al*. (2017), Asmamaw Amogne *et al*. (2020), Wakjira Getachew *et al*. (2021), and Desalegn Nigatu *et al. (*2022) justify a positive correlation between hundred seed weight with branches per plant, contrasting with these findings, a negative correlation of hundred seed weight with days to maturity, seed per pod, pod per plant, and plant height at phenotypic level was reported by Chaudhary *et al*. (2020).

The aboveground dry biological yield (biomass yield) had a highly significant positive phenotypic correlation with the seed yield (rp=0.44**) and a positive significance correlation with the number of branches per plant (0.19*), while a highly significant negative phenotypic correlation with the harvest index (rp= −0.34**) indicated that an increase in the biological yield will help to increase the seed yield of common bean simultaneously. A negative correlation of biological yield with harvest index implies difficulty in improving both traits at the same time through direct selection or the two traits are governed by the same physiological pathway. In line with this, Kassaye Negash (2006) and Itefa Degefa *et al*. (2014) and found a highly significant positive correlation between biological yield and seed yield at the phenotypic level whereas Kedir Gelgelu *et al*. (2017) reported a high significance negative phenotypic correlation between biological yield and harvest index. Harvest index had a highly significant positive phenotypic correlation with seed yield (rp = 0.67**) while a highly significant negative phenotypic correlation with above-ground biomass (rp= −0.34**) and a non-significant phenotypic correlation with other traits included in this study. Similarly, a positive significance of harvest index with biological yield and highly positive significance with seed yield were reported by (Itefa Degefa *et al.,* 2014; Desalegn Nigatu *et al.,* 2022, and Kedir Yimam *et al.,* 2023). However, Kassa Mammo *et al*. (2019) reported a positive phenotypic correlation between harvest indexes with biological that disagrees with these findings.

#### Genotypic correlations of seed yield with other traits

The estimated genotypic correlation coefficients between seed yield with other possible pairs of seed yield and related traits of common bean genotypes are presented in Table 3.

Genotypic correlation coefficients between seed yield and seed yield-related traits ranged from −0.27 to 0.74. In this study, seed yield had highly significant positive genotypic correlations with above-ground biomass) (rg= 0.51**), harvest index (rg=0.74**) and significant genotypic correlation with branches per plant (0.197*). The positive correlation of branch per plant, biological yield, and harvest index with seed yield might indicate the presence of strong linkage of genes or the characters may be the result of pleiotropic genes that control these characters in the same direction. This suggests a common genetic/physiological basis among these traits and the possibility of simultaneous improvement of these traits.

The results of this study demonstrate that genotypes with more branches per plant, maximum biomass, and harvest index would produce a higher seed yield than those with lower levels of these traits. Selection of genotypes based on these traits would be effective in increasing seed yield. In line with the present study conducted by Chaudhary *et al*. (2020), Desalegn Nigatu *et al*. (2022), Welde Ketema and Negash Geleta (2022), and Belay Asmare *et al*. (2023) reported a positively and significantly correlated branch per plant, biological yield and harvest index with seed yield at the genotypic level.

Conversely, seed yield exhibited a negative significance genotypic correlation with days to maturity (rg= −0.31*) and the negative genotypic correlation obtained between days to maturity with seed yield indicates that common bean seed yield can be improved by selecting early maturing genotypes in the moisture stress areas. Other findings done by Tesfaye Walle *et al*. (2018), Chaudhary *et al*. (2020) and Belay Asmare *et al*. (2023) report a negative genotypic correlation of days to maturity with seed yield on cowpea and common bean genotypes respectively. But in contrast with this, Desalegn Nigatu *et al*. (2022) report a non-significance correlation between days to maturity and seed yield at the genotypic level. Days to flowering had a negatively significant genotypic correlations with branch per plant (rg= −0.34*) and non-significance with the reaming traits included in the study (Table 3). This negative significance correlation at the genotypic level indicates that the two traits may either linked or follow the same physiological pathway and genotypes that flowering early may not produce more branches.

Days to maturity had negative highly significant genotypic correlation with plant height (rg=-0.5**), branches per plant (rg= −0.34**), and significant negative correlation at genotypic level with seed per pod (rg=-0.27*), seed yield (rg= −0.31*) and harvest index (rg = −0.29*) and with the rest of traits, there was no significant correlation at the genotypic level. Plant height had a highly significant negative genotypic correlation with days to maturity (rg= −0.5**) and non-significant correlation with the remaining traits in this study. Branches per plant had a highly positive and significant correlation with hundred seed weight (rg= 0.33**) and a positive significant correlation with pods per plant (rg=0.29*), seeds per pod (rg= 0.26*), and aboveground biological yield (rg= 0.27*) at the genotypic level but have highly negative significant genotypic correlations with days to maturity (rg= −0.34**) and negative significant genotypic correlations with days to flowering) (rg= −0.3*) while with the rest, there was no significant correlation. Pod per plant had significant positive genotypic correlations with the number of branches per plant (rg=0.29*) and with 100-seed weight (rg=0.31*) at the genotypic level, while non-significant with other studied characters under the study.

Seed per pod had a significant positive genotypic correlation with branches per plant (rg= 0.26*) and negative significance (rg= −0.27*) with days to maturity. However, it was non-significant with other traits. The negative association of seeds per pod with maturity date suggested that the early maturity genotype may not have maximum seed per pod and improving such traits simultaneously through direct means is impractical. Hundred seed weight had a highly significant positive genotypic correlation with branches per plant (rg=0.33**) and a significant positive correlation with pods per plant (rg=0.31*), while it had a significant negative correlation with the harvest index (rg= −0.32*) and non-significant with other traits in this study. In line with these findings, genotypic positive correlation of hundred seed weight with branch per plant and pod per plant was done by Belay Asmare *et al*. (2023). But contrasting with this, a negative significant genotypic correlation of hundred seed weight with pod per plant was done by Kedir Yimam *et al.,* 2023).

Aboveground biomass had a highly significant positive correlation with seed yield (rg=0.51**) and a significant positive correlation with branch per plant (rg=0.27 *) at the genotypic level. There was non-significant correlation with days to flowering, days to maturity, plant height, seed per pod, pod per plant, or harvest index (Table 3). Also, harvest index shows a highly significant positive correlation with seed yield (rg= 0.74**) and a significant negative correlation with days to maturity (rg= −0.29*) and hundred seed weight (rg= −0.32*) at the genotype level.

In this study, the magnitudes of genotypic correlation coefficients were greater than the corresponding phenotypic correlation coefficients for most of the study traits. This indicates that the correlation was more due to genotypic and phenotypic correlations being slightly influenced by environmental effects. However, the direction of the correlation was the same at both genotypic and phenotypic levels. Similarly, a higher genotypic correlation coefficient than phenotypic correlation coefficient was found for most of the traits by (Mesfin Hailemariam, 2018; Sima Jibril *et al.,* 2019; Simon Yohannes *et al.,* 2020; Belay Asmare *et al.,* 2023 and Kedir Yimam *et al.,* 2023).

Based on this correlation result, seed yield had a significant and positive correlation both at genotypic and phenotypic level with branches per plant, aboveground biomass, and, harvest index, implying the possibility of a correlated response to selection. Therefore, these traits could be used as indirect selection criteria during genotype evaluation of common bean in the study area.

### Path coefficient analysis

#### Phenotypic path analysis

The path analysis included characters that had a significant correlation (both negative and positive) with seed yield (Dewy and Lu, 1959). Path coefficients were classified as negligible (0.00 - 0.09), low (0.10 - 0.19), moderate (0.20 - 0.29) or high (0.30 - 0.99), with values above 1.00 considered very high (Lenka and Misra, 1973). According to this description, the harvest index (0.940) and biological yield (0.748) had significant positive direct effects on seed yield (Table 4). They also showed a strong positive correlation with seed yield (Table 4). High direct effects indicate a real relationship between these traits and seed yield. Therefore, selection for these traits would have a reasonable effect and improve seed yield in common bean. These traits should be considered as important selection criteria in common bean breeding programme, as an increase in these traits will lead to an increase in seed yield. According to Awol Mohammed and Asnake Fikre (2018), Kassa Mammo *et al*. (2019), Amare Tsehaye *et al*. (2020) and Kedir Yimam *et al*. (2023), the greatest positive direct effect on seed yield was observed for the biological yield and harvest index.

**Table 4.** Estimates of direct (bold) and indirect effects (off the diagonal) of different traits on seed yield at the phenotypic level in 64 small-seeded common bean genotypes.

| Traits | Df | Dm | PH | Bpp | PPP | Spp | Hsw | AGB | HI | Rg |
| --- | --- | --- | --- | --- | --- | --- | --- | --- | --- | --- |
| Df | **-0.017** | -0.003 | -0.004 | 0.0011 | 0.0009 | 0.0047 | -0.007 | -0.090 | -0.752 | 0.090 |
| Dm | -0.002 | **-0.026** | -0.006 | 0.0008 | -0.008 | 0.0021 | -0.008 | -0.052 | -0.315 | 0.047 |
| PH | 0.002 | 0.0006 | **0.034** | -0.0007 | -0.009 | -0.026 | 0.022 | 0.016 | 0.1324 | 0.170 |
| Bpp | 0.005 | 0.0005 | 0.006 | **-0.004** | -0.017 | -0.083 | 0.036 | 0.1442 | 0.0520 | 0.190 |
| PPP | 0.002 | -0.003 | 0.004 | -0.009 | **-0.074** | -0.017 | 0.035 | -0.114 | 0.0909 | 0.070 |
| Spp | 0.003 | 0.0002 | 0.003 | -0.011 | -0.004 | **-0.030** | 0.004 | -0.417 | 0.1394 | 0.070 |
| Hsw | 0.001 | 0.0002 | 0.006 | -0.012 | -0.021 | -0.009 | **0.012** | 0.0789 | -0.153 | 0.060 |
| AGB | 0.002 | 0.0000 | 0.001 | -0.008 | 0.0001 | 0.0017 | 0.013 | **0.7461** | -0.315 | 0.440 |
| HI | 0.002 | 0.0001 | 0.048 | -0.002 | -0.007 | -0.044 | -0.020 | -0.250 | **0.9411** | 0.670 |

Residual effect = 0.027. Note: BPP = branch per plant, AGB = aboveground biological yield per hectare, HI= harvest index, and SyrP = phenotypic correlation coefficient with seed yield.

At the genotypic level, number of branches per plant had a negligible direct effect on seed yield (0.00017). The phenotypic residual effects (R=0.027) were low, indicating that 97.3% of the seed yield of common bean was determined by the traits studied in this experiment. Other independent variables or seed yield related traits that are not included in this experiment are expected to have a small effect of only 2.7% on seed yield. This result suggests that the traits included in this study are adequate, although additional traits may be needed to fully explain seed yield. This finding is consistent with Meseret Tola *et al*. (2020) who reported that 92.9% of the variation in seed yield in common bean genotypes at the phenotypic level was due to causal factors, while only 7.1% was due to other factors that were not included in their study. Similarly, Belay Asmare *et al*. (2023) found that 91.8% of the variation in seed yield was due to causal factors and only 8.2% was due to other factors not included in their study at the phenotypic level in common bean genotypes. A lower R value than that of the cited authors may be due to environmental effects or to fewer or more parameters collected during the study.

#### Genotypic path analysis

Table 5 presents the genotypic direct and indirect effects of various traits on seed yield. At the genotypic level, path coefficient analysis revealed that the harvest index (0.85) and aboveground biological yield (0.67) had positive direct effects on seed yield, indicating that these yield components positively influenced seed yield in common bean. Indirect selection based on the harvest index and biological yield can effectively improve seed yield in small-seeded common bean. Studies by Kassa Mammo *et al*. (2019) and Kedir Yimam *et al*. (2023) have shown that these traits have a strong positive direct effect on seed yield at the genotypic level, indicating a true relationship. However, Desalegn Nigatu *et al*. (2022) report that the harvest index has an indirect effect on seed yield.

**Table 5.**
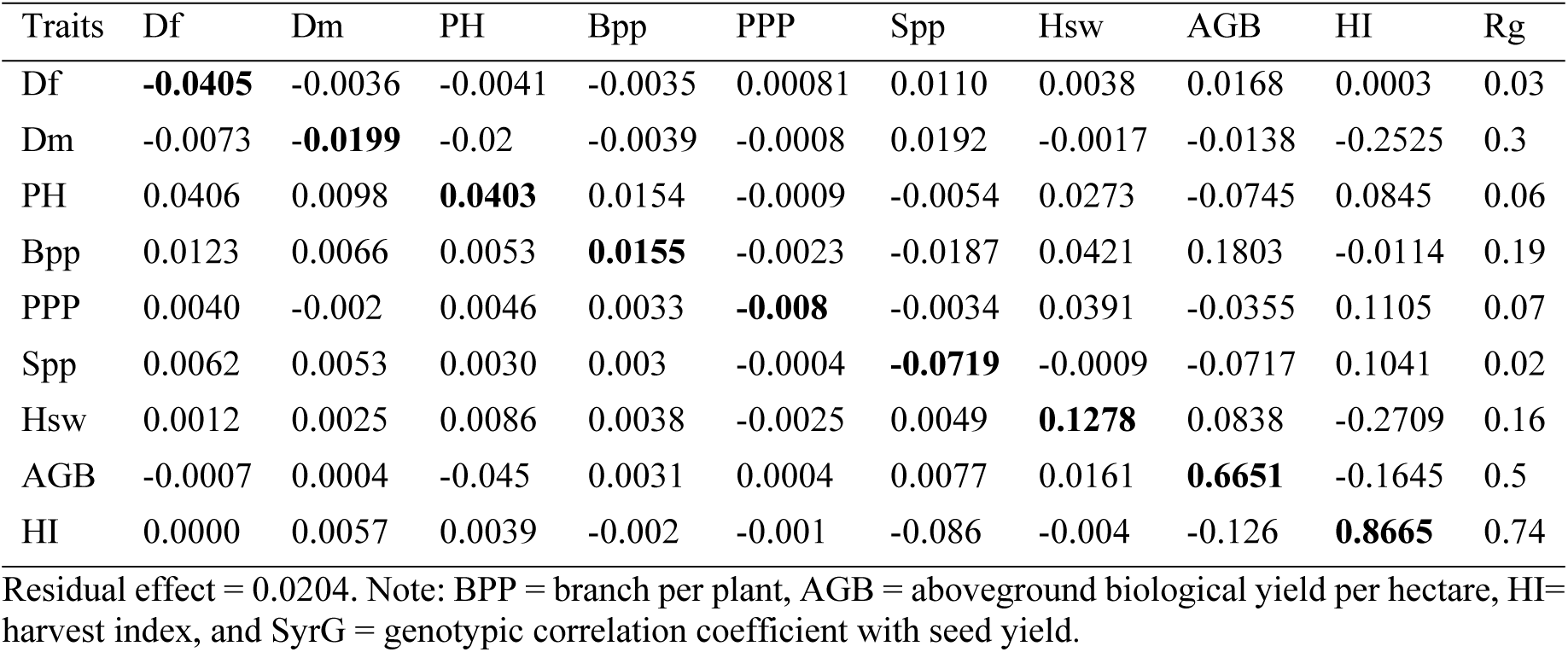
Estimates of direct (bold) and indirect effects (off the diagonal) of different traits on seed yield at the genotypic level in 64 small-seeded common bean genotypes.

| Traits | Df | Dm | PH | Bpp | PPP | Spp | Hsw | AGB | HI | Rg |
| --- | --- | --- | --- | --- | --- | --- | --- | --- | --- | --- |
| Df | **-0.0405** | -0.0036 | -0.0041 | -0.0035 | 0.00081 | 0.0110 | 0.0038 | 0.0168 | 0.0003 | 0.03 |
| Dm | -0.0073 | **-0.0199** | -0.02 | -0.0039 | -0.0008 | 0.0192 | -0.0017 | -0.0138 | -0.2525 | 0.3 |
| PH | 0.0406 | 0.0098 | **0.0403** | 0.0154 | -0.0009 | -0.0054 | 0.0273 | -0.0745 | 0.0845 | 0.06 |
| Bpp | 0.0123 | 0.0066 | 0.0053 | **0.0155** | -0.0023 | -0.0187 | 0.0421 | 0.1803 | -0.0114 | 0.19 |
| PPP | 0.0040 | -0.002 | 0.0046 | 0.0033 | **-0.008** | -0.0034 | 0.0391 | -0.0355 | 0.1105 | 0.07 |
| Spp | 0.0062 | 0.0053 | 0.0030 | 0.003 | -0.0004 | **-0.0719** | -0.0009 | -0.0717 | 0.1041 | 0.02 |
| Hsw | 0.0012 | 0.0025 | 0.0086 | 0.0038 | -0.0025 | 0.0049 | **0.1278** | 0.0838 | -0.2709 | 0.16 |
| AGB | -0.0007 | 0.0004 | -0.045 | 0.0031 | 0.0004 | 0.0077 | 0.0161 | **0.6651** | -0.1645 | 0.5 |
| HI | 0.0000 | 0.0057 | 0.0039 | -0.002 | -0.001 | -0.086 | -0.004 | -0.126 | **0.8665** | 0.74 |

Residual effect = 0.0204. Note: BPP = branch per plant, AGB = aboveground biological yield per hectare, HI= harvest index, and SyrG = genotypic correlation coefficient with seed yield.

In addition, days to maturity (−0.038) had a negative direct effect on seed yield at the genotypic level (Table 5), likely due to its negative correlation with seed yield at the genotypic level (Table 5). These negative effects may lead to a decrease in seed yield. The data shows that early maturing accessions had a higher seed yield per hectare than late-maturing accessions. Therefore, selecting against this trait could increase the seed yield per hectare. This result is consistent with the findings of Ejigu Ejara *et al*. (2017) and Kedir Shafi *et al*. (2019), who reported a negative direct effect of days to maturity on the seed yield of common bean genotypes. However, Tesfaye Walle *et al*. (2018) and Meseret Tola *et al*. (2020) reported a positive correlation between days to maturity and seed yield. This may be because under optimal environmental conditions, a longer time to maturity increases the seed yield, likely because more photosynthesis is diverted to the seed. Singh and Chaudhary (1977) suggested that when traits have a positive correlation and strong positive indirect effects but negative direct effects on economic traits such as seed yield, emphasis should be given to indirect effects. Therefore, traits that are strongly and truly associated with grain yield can be used as indirect selection criteria. Biomass yield and the harvest index can be used as indirect selection criteria for improving common bean seed yield in the area.

The genotypic residual effect (R=0.02) showed that 98% of the seed yield of common bean was contributed by the studied traits in this path analysis. The experiment expected that other independent variables not included would only have a 2% influence on seed yield. This result shows that the traits included in this path analysis study are adequate, even though other traits may also be necessary. Similarly, Belay Asmare *et al*. (2023) reported that 97.39% and 95.99% of the total variation in seed yield was explained by the study variables at the genotypic level for common bean genotypes in Goro and Ginnir, Southeast Ethiopia. In line with this result, Desalegn Nigatu *et al*. (2022) reported that 94% of the variability in seed yield was explained by the study variables at genotypic levels on soybean genotypes at Pawe.

### Cluster Analysis and Genetic Divergence

#### Cluster analysis

In this study, the results of the pseudo F test showed that eight non overlapping cluster arrangements were the most suitable for grouping all 64 genotypes, including the standard checks. The genotypes were then grouped into eight distinct clusters, forming different hierarchical subgroups (Figure 2 and Table 6). Similar reported done by Belay Asmare *et al*. (2023) grouped 64 small-seeded common bean genotypes into eight genetically distinct clusters in the Bale and East Bale Zones. Similarly, Eyuel Mesera *et al*. (2022) grouped 100 common bean genotypes into eight clusters. Lima *et al*. (2012) also grouped 100 common bean genotypes into eight groups in Brazil. These findings suggest that the tested genotypes are genetically diverse and have different backgrounds. Cluster I had the highest number of genotypes, with 27, while cluster VIII had the lowest, with only three genotypes. A total of 26 new genotypes and one standard check variety constituted for 42.18%, were grouped into cluster I, whereas cluster VIII contained only 3 genotypes, accounting for 12.5%. Cluster V comprised 8 genotypes (10.93%), including seven new genotypes and one standard check. Cluster VII contained 7 genotypes (10.93%), including 6 new genotypes and one standard check variety. Cluster II include six new genotypes (9.37%), while cluster IV included 5 new genotypes (7.8%). Both cluster III and cluster VI contained 4 (6.25%) new genotypes each (Table 6). The standard check variety (genotypes 62, 63, and 64) was grouped in clusters I, V, and VII, indicating a close relationship with these clusters compared to other genotype clusters.

**Figure 2.**
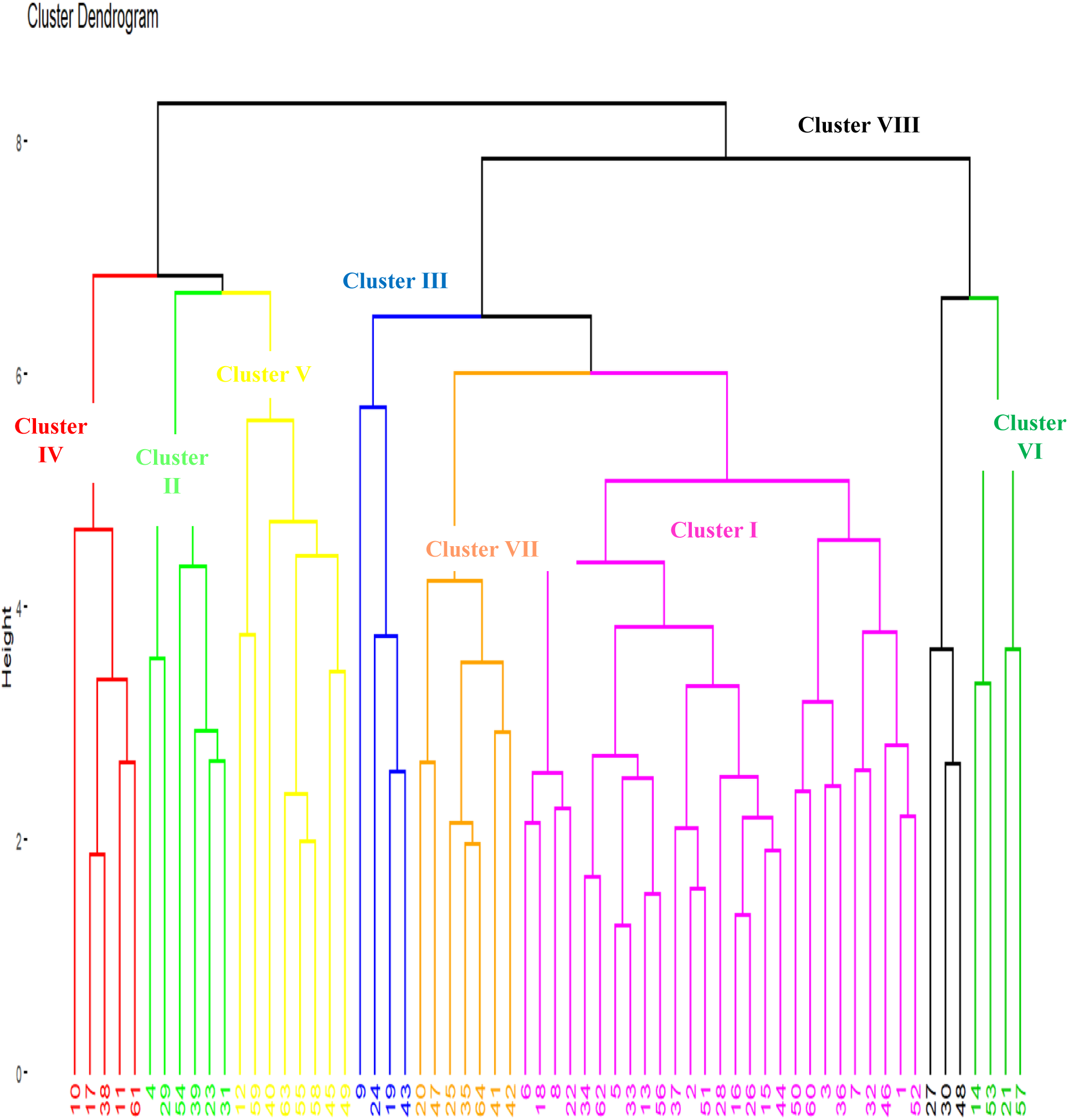
Dendrogram constructed using 10 traits of 64 common bean genotypes used for the study.

**Table 6.** Distribution and grouping of 64 small-seeded common bean genotypes into eight different clusters based on the Mahalanobis (D2) distance for 10 traits.

| Cluster | No genotype | proportion in % | List of Genotypes |
| --- | --- | --- | --- |
| I | 27 | 42.18 | 6, 18, 8, 22, 34, 62, 5, 33, 13, 56, 37, 2, 51, 28, 16, 26, 15, 44, 50, 60, 3, 36, 7, 32, 46, 1, 52 |
| II | 6 | 9.37 | 4, 29, 54, 39, 23, 31 |
| III | 4 | 6.25 | 9, 24, 19, 43 |
| IV | 5 | 7.83 | 10, 17, 38, 11, 61 |
| V | 8 | 12.5 | 12, 59, 40, 63, 55, 58, 45, 49 |
| VI | 4 | 6.25 | 14, 53, 21, 57 |
| VII | 7 | 10.94 | 20, 47, 26, 35, 64, 41, 42 |
| VIII | 3 | 4.68 | 27, 30, 48 |

#### Cluster means analysis

The mean values of the 10 quantitative traits for each cluster. Compared with the other clusters, the cluster I genotypes were characterized by longer flowering times, shorter plant heights, greater numbers of branches per plant, greater aboveground biomass, lower seed yields, and lower harvest index (Table 7). Cluster II genotype had a medium seed yield, tallest plant height, greater number of branches per plant, greater number of seeds per pod, medium hundred seed weight, and medium biomass values, higher seed yield and higher harvest index. The cluster III genotypes had longer days to flowering and maturity, shorter plant heights, lower numbers of branches, medium numbers of pods per plant, high to medium aboveground biomass, lower hundred seed weights, greater seed yields, and higher harvest indices.

**Table 7.** Mean values of clusters for 10 traits of 64 common bean genotypes.

| Clusters | DsF | Dsm | Ph | Bpp | PPP | Spp | HsW | AGB | SY | HI |
| --- | --- | --- | --- | --- | --- | --- | --- | --- | --- | --- |
| C1 | 54.9 | 84.1 | 35.62 | 4.48 | 15.92 | 4.61 | 29.05 | 7504.02 | 2277.76 | 30.97 |
| C2 | 53.47 | 82.18 | 44.51 | 4.79 | 17.58 | 4.64 | 27.95 | 6610.68 | 2699.54 | 41.19 |
| C3 | 56.07 | 84.29 | 37.11 | 3.09 | 15.41 | 4.24 | 23.17 | 6562.36 | 2912.17 | 45.24 |
| C4 | 53.4 | 83.1 | 42.76 | 5.2 | 22.2 | 3.78 | 32.74 | 7654.8 | 3060.94 | 40.29 |
| C5 | 53.19 | 76.25 | 48.54 | 4.53 | 14.11 | 4.53 | 30.24 | 7011.29 | 2873.31 | 40.23 |
| C6 | 56.29 | 83.93 | 41.74 | 3.84 | 13.8 | 3.65 | 25.56 | 6292.2 | 2103 | 34.12 |
| C7 | 54.33 | 83.33 | 49.83 | 4.27 | 27.5 | 4.33 | 33.9 | 6656.83 | 2321.13 | 34.98 |
| C8 | 54.86 | 81.71 | 46.1 | 3.79 | 16.11 | 4.37 | 31.47 | 5574.32 | 1956.2 | 35.11 |

Where:- c1-c8 are number of clusters, Dsf days to flowering, Dsm, days to maturity, Ph= plant height, Bpp= number of braches per plant, Ppp= number of pods per plant, Spp= number of seeds per pod, Hsw= 100 seed weight, AGB= aboveground biomass, Sy= seed yield and HI= harvest index

Cluster IV genotype is characterized by a longer time to flowering and maturity, shorter plant height, medium to high number of branches per plant, and low to high number of pods per plant. It also has the maximum number of seeds per pod, medium to high hundred seeds weight, higher aboveground biomass, low seed yield, and low harvest index compared to those of the other clusters. The cluster V genotype exhibited shorter days to flowering and maturity, taller plant heights, fewer pods per plant, greater aboveground biomass, greater seed yield, and greater harvest index. The plants of the cluster VI genotype, on the other hand, had fewer pods per plant and fewer seeds per pod, medium aboveground biomass, lower seed yield, and a lower harvest index. Cluster VII genotypes were distinguished by longer days to maturity, tallest plant height, maximum number of pods per plant, medium number of branches per plant, higher hundred seed weight, medium aboveground biomass, lower seed yield and harvest index. Cluster VIII genotypes were characterized by late flowering and maturity, lower aboveground biomass, greater hundred-seed weight, lower seed yield, and medium harvest index.

In this study, clusters III, IV, and V were identified as high yielders, while clusters VII and VIII were identified as low yielders.

By crossing clusters with wider inter cluster distances that have superior mean performance for desirable traits, reasonable changes can be achieved. Cluster V was found to be early maturing. Cluster IV was identified as a high yielder, and cluster V was identified as both early maturing and high yielding. In contrast, clusters VII and VIII were identified as late maturing clusters with low seed yield traits. Therefore, crossing clusters IV and VII will result in a new high-yielding genotype. Similarly, crossing clusters V and VIII will produce new genotypes that are both early maturing and high yielding, making them suitable for moisture deficit environments such as the study areas due to their wider genetic distance.

#### Genetic divergence analysis

The genotypes were categorized into eight distinct clusters based on a cluster analysis of 10 quantitative traits. The chi-square test showed statistically significant differences between most of the clusters. Table 8 presents the average intra cluster (differences among genotypes within the same cluster) and inter cluster (differences among genotypes between different clusters) distance (D2) values. The inter-cluster distances ranged from 10.21 to 351.39, while the intra-cluster distances ranged from 2.96 to 4.22 (Table 8). The results show that the distances between clusters were greater than those within clusters, indicating variability among the tested genotypes of different groups and both heterogeneous and homogeneous nature within and between clusters.

**Table 8.** Average intra (bold face) and inter pairwise generalized square distance (D2) values of eight clusters of 64 common bean genotypes.

| Clusters | C1 | C2 | C3 | C4 | C5 | C6 | C7 | C8 |
| --- | --- | --- | --- | --- | --- | --- | --- | --- |
| C1 | **4.22** | 10.21<sup>ns</sup> | 43.79** | 96.56** | 20.23* | 60.43** | 35.94** | 201.22** |
| C2 |  | **3.29** | 13.97<sup>ns</sup> | 48.86** | 52.03** | 29.35** | 76.05** | 128.12** |
| C3 |  |  | **2.96** | 19.45* | 111.78** | 18.38* | 131.45** | 70.25** |
| C4 |  |  |  | **3.62** | 196.79** | 57.06** | 204.54** | 38.79** |
| C5 |  |  |  |  | **3.04** | 117.03** | 33.34** | 331.13** |
| C6 |  |  |  |  |  | **3.2** | 149.99** | 92.78** |
| C7 |  |  |  |  |  |  | **3.99** | 351.39** |
| C8 |  |  |  |  |  |  |  | **3.38** |

Where ns= non-significant and \*, \*\* significance at the 5% and significance at the 1% probability level, and $X^2 = 21.67$ at 5% and 16.92 at 1%.

The largest inter-cluster distance was observed between clusters VII and VIII (D2 = 351.39), followed by clusters V and VIII (D2 = 331.23), IV and VIII (D2 = 204.54), IV and V (D2 = 196.79), and VI and VII (D2 = 149.99), indicating greater genetic divergence between these clusters. The genotype members of distant clusters (clusters VII and VIII, V and VIII, IV and VII, IV and V, and clusters VI and VII) can be used in breeding programmes to achieve a high level of heterotic expression in the F1 generation. This can be achieved by combining desirable traits with greater heterotic potential and a wider range of variability in the recombinant and segregating F2 generation. The selection of better parents from these clusters for hybridization programmes can help to achieve ideal segregates and/or recombinants. The minimum distance between clusters was recorded between clusters I and II (D2 = 10.21), followed by clusters II and 3 (D2 = 13.97), clusters III and VI (D2 = 18.38), and clusters III and IV (D2 = 19.45). The close genetic relationship between genotype members in these clusters suggests that crossing genotypes from these clusters may not result in greater heterotic value in F1 and a wide range of variability in the segregating F2 population. Cluster 1 had the greatest average intracluster distance (D2 = 4.22), indicating greater genetic divergence among its members than among the other clusters. Cluster III had the lowest intracluster distance (D2 = 2.96), followed by cluster 5 (D2 = 3.04). Similarly, genetic divergence was reported in common bean genotypes based on clustering analysis by Nigussie Kefelegn *et al*. (2020), Eyuel Mesera *et al*. (2022), Kedir Yimam *et al*. (2023), and Belay Asmare *et al*. (2023).

### Principal Component Analysis

The genotypes tested showed genotypic variability, as confirmed by principal component analysis. The results of principal component analysis for 10 traits evaluated in 64 small-seeded common bean genotypes are presented in table 9. The percentages of total variance ranged from 10.5% to 21.4%. The first five principal component axes (PCA1 to PCA5) accounted for 74.3% of the total variation, with eigenvalues greater than unity (Table 9). The remaining 25.7% was explained by the last five PCs with eigenvalues less than unity, indicating variability in the genotypes for the studied traits. The cumulative contribution of these three components is 52.1%, which accounts for more than half of the total variation. Abdi and Williams (2010) state that the first principal component has the highest variance and explains the largest part of the variability. The first PC explains 21.4% of the variation, and was the highest among all the PCs, followed by PC2 (16.9%), PC3 (13.8%), PC4 (11.7%), and PC5 (10.5%) (Table 9). PC1 was positively loaded and exhibited relatively high variation in terms of seed yield, branch per plant, harvest index, and plant height. The second principal component (PC2) varied mainly due to the seed yield and harvest index, which had a positive loading, and the hundred seed weight and branches per plant, which had a negative loading. The traits that contributed the most to the variation in PC3 were a negative loading of aboveground biomass and seed yield, while the number of seeds per pod had a positive loading. The variation in the fourth principal component was explained more by the number of days to maturity and the number of pods per plant, both of which had positive loadings. The fifth principal component was mostly associated with the number of seeds per pod, with a positive loading and was negatively associated with the number of days to flowering, plant height, and hundred-seed weight. Thus, the traits that exhibited large contribution to the genetic variability of common bean genotypes. This is consistent with the findings of Nadeem *et al*. (2020), who reported that the first five PCs explained 71% of the total variation. Esrael Kanbata (2023) reported that the first five principal components explained 71.88% of the total variation, with PCA1, PCA2, and PCA3 contributing 21.27%, 16.21%, and 13.24%, respectively.

**Table 9.** Eigenvectors, eigenvalues, percentages and cumulative variances of the first five principal components (PCs) for 10 traits of 64 common bean genotypes.

| Trait | PC1 | PC2 | PC3 | PC4 | PC5 |
| --- | --- | --- | --- | --- | --- |
| Days to 50% flowering | 0.298 | 0.104 | 0.181 | 0.096 | -0.415 |
| Days to 90% maturity | -0.202 | 0.080 | -0.106 | 0.692 | 0.257 |
| Plant height | 0.331 | -0.064 | 0.142 | -0.213 | -0.529 |
| Branch per plant | 0.454 | -0.300 | 0.010 | -0.045 | 0.226 |
| Pod per plant | 0.266 | -0.210 | 0.162 | 0.614 | -0.150 |
| Seed per pod | 0.272 | -0.019 | 0.304 | -0.104 | 0.534 |
| 100- seed weight | 0.209 | -0.506 | 0.041 | 0.209 | -0.301 |
| Above ground biomass | 0.158 | -0.220 | -0.777 | -0.070 | 0.128 |
| Harvest index | 0.354 | 0.610 | 0.171 | 0.150 | -0.117 |
| Seed yield | 0.464 | 0.410 | -0.428 | 0.082 | -0.045 |
| Eigen value | 2.130 | 1.69 | 1.37 | 1.17 | 1.04 |
| Proportion variance in each PC (%) | 21.4 | 16.9 | 13.8 | 11.7 | 10.5 |
| Cumulative variance (%) | 21.4 | 38.3 | 52.1 | 63.8 | 74.3 |

PC=Principal component

## CONCLUSIONS AND RECOMMENDATIONS

The analysis of variance revealed highly significant (P<0.01) differences among the genotypes for most traits considered and indicated the existence of variation among the tested genotypes. The highest seed yield was recorded for genotype number 9 (3702.10 kg ha^-1^), while the lowest was obtained for genotype number 63 (1633.1 kg ha^-1^). Compared with the three check varieties (genotype numbers 62, 63, and 64) the genotypes had a seed yield advantage of 24.56% to 55.89%. The plant height, number of branches per plant, hundred-seed weight, biomass, seed yield, and harvest index were had moderate genotypic coefficients of variation. High heritability associated with a high genetic advance as a percentage of the mean (GAM) was found for plant height, number of branches per plant, number of pods per plant, and hundred-seed weight, whereas seed yield showed moderate heritability with a high GAM.

Seed yield was positively and significantly correlated with the number of branches per plant, aboveground biomass, and harvest index, and negatively and significantly correlated with the number of days to maturity at both the genotypic and phenotypic levels. Path coefficient analysis revealed that the number of branches per plant, aboveground biomass, and harvest index had direct effects on seed yield. The cluster analysis grouped the genotypes into eight distinct clusters. The greatest intercluster distance was found between cluster VII and VIII, followed by cluster V and cluster VIII, as well as between cluster IV and cluster VII. The first five PCs with eigenvalues greater than one explained 74.3% of the total variation in which days to maturity, plant height, pods per plant, seed per pod, seed yield, and harvest index were the traits that contributed most to the variation. Overall, the results of this study indicated the presence of genetic variability and trait associations among the 64 tested common bean genotypes as well as the possibility of designing effective genetic improvement methods. It is recommended that genotypes that have high agronomic performance (early days to flowering and days to maturity and high seed yield) could be included in further breeding stages for common bean improvements in the study area. For reliable results and recommendations, it is better to repeat the experiment for at least one additional season across locations for the development of high yielder and early maturing common bean genotypes.

## Accessibility of the data (Data Availability)

Data used to support the findings of this study are available from the first author upon request.

## Acknowledgments

Heartily we acknowledge to Melkassa Agricultural Research Center for the aid of planting materials (seed) and would like to thank Mr. Kindalm Yargal, Mr. Fiker kebede, Mr. Gizachaw Haile Maryam, and Mr. Alemu Lakew for their technical support at field level up to the end of this study.

## Appendix

**Appendix Table 1.** List and description of small seeded common bean genotypes (materials) used for the experiment.

| Genotype number | Genotype Code | Genotype number | Genotype code |
| --- | --- | --- | --- |
| 1 | CB170000194 | 34 | CB170003-16 |
| 2 | Geno#126 | 35 | CB1700037-28-2 |
| 3 | CB1700029.11 | 36 | CB17000383-4-1 |
| 4 | CB170023-5-2 | 37 | CB170038-11-1 |
| 5 | CB170065-59-3 | 38 | CB170008-4 |
| 6 | CB170064102 | 39 | CB17100040321 |
| 7 | CB17000319 | 40 | CB17005232 |
| 8 | CB170023-4 | 41 | GENO206 |
| 9 | CB18100254-1 | 42 | CB170001-18 |
| 10 | CB1700254 | 43 | GENO 126 |
| 11 | CB17000320 | 44 | GENO 331 |
| 12 | CB1700081 | 45 | CB181000862 |
| 13 | CB1700025-23 | 46 | CB170065591 |
| 14 | CB17004612 | 47 | CB170005824 |
| 15 | CB1700254 | 48 | CB170052292 |
| 16 | CB1700085 | 49 | CB17001535-3 |
| 17 | CB17000253 | 50 | CB170052292 |
| 18 | CB170065.56 | 51 | GENO161 |
| 19 | Geno34 | 52 | CB17003231 |
| 20 | GENO 418 | 53 | CB176685593 |
| 21 | CB170058122 | 54 | CB24223 |
| 22 | GENO214 | 55 | CB170003317 |
| 23 | CB170038-15 | 56 | CB17000320 |
| 24 | CB1810004032-3 | 57 | CB17002952 |
| 25 | CB17000324-1 | 58 | CB17000317 |
| 26 | CB170003-20 | 59 | CB1700029141 |
| 27 | CB170002-10 | 60 | CB170006210 |
| 28 | B12262 | 61 | CB17003619 |
| 29 | CB1700037-18 | 62 | AWASH -2 |
| 30 | GENO147 | 63 | Awash Miten |
| 31 | CB170003-241 | 64 | Awash Melka |
| 32 | CB170002-11-1 |  |  |
| 33 | CB170003-16 |  |  |

Source: Melkassa Agricultural Research Center (MARC), Low land pulses Breeding Program (2023)

**Appendix Table 2.**
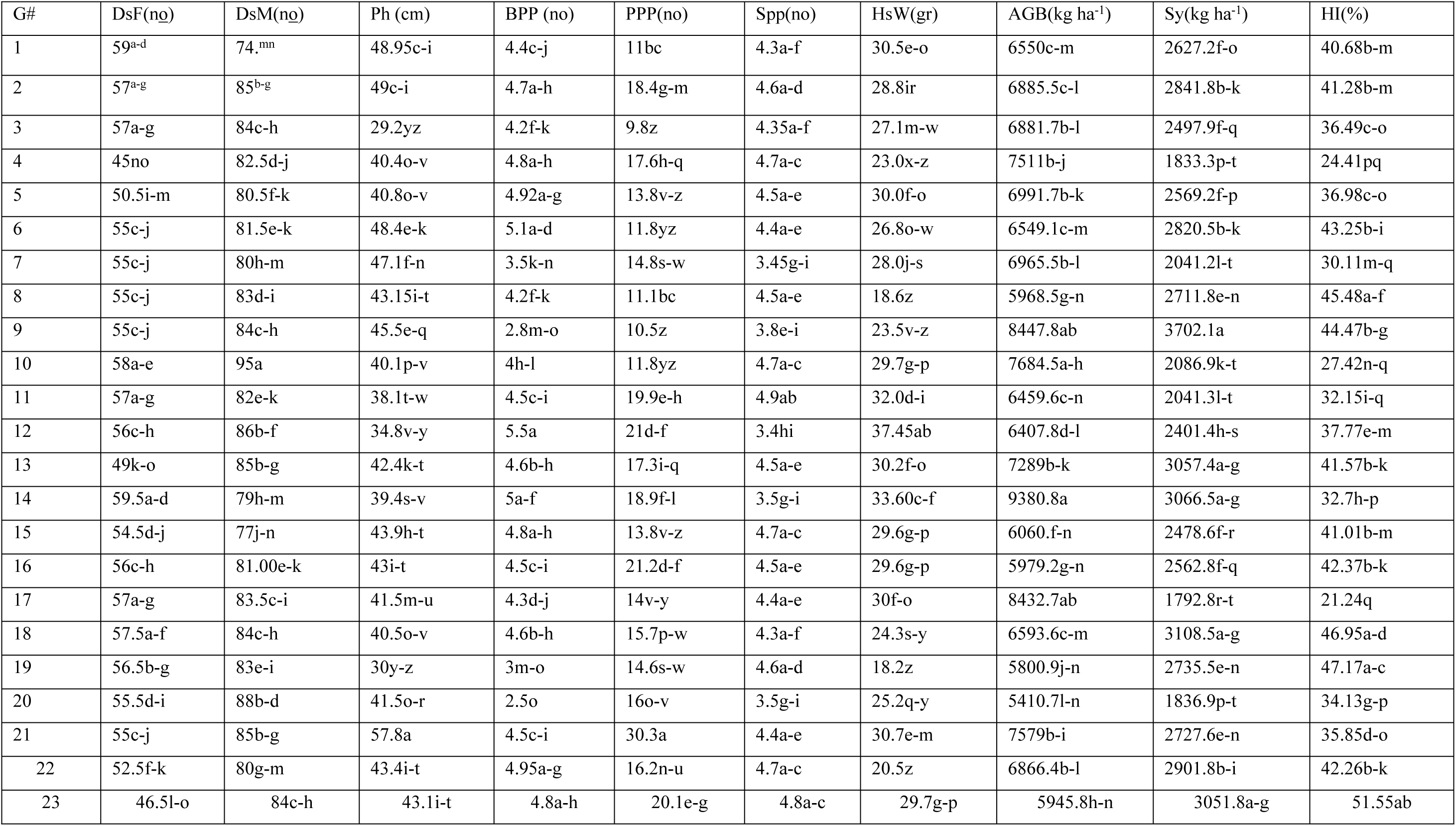
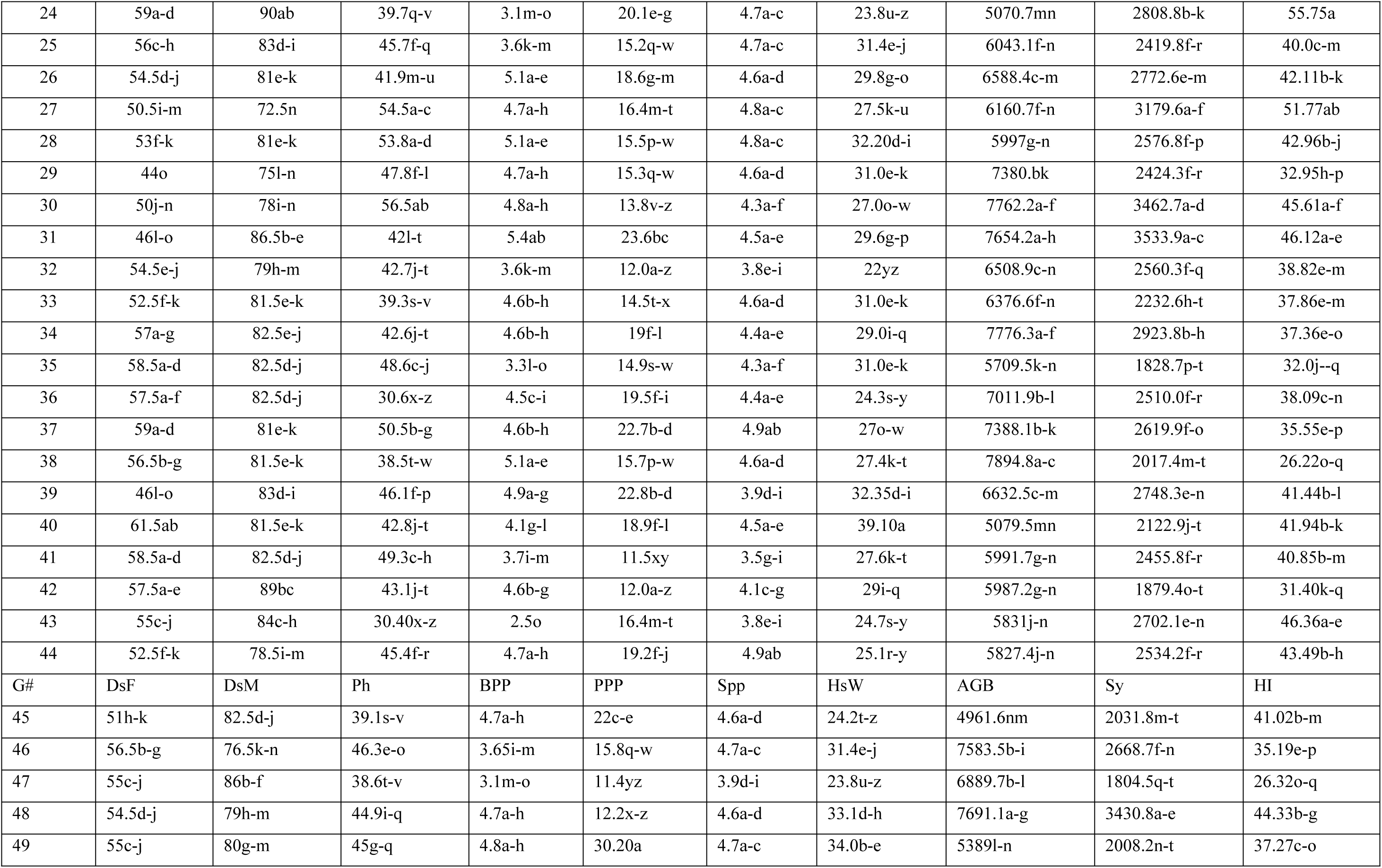
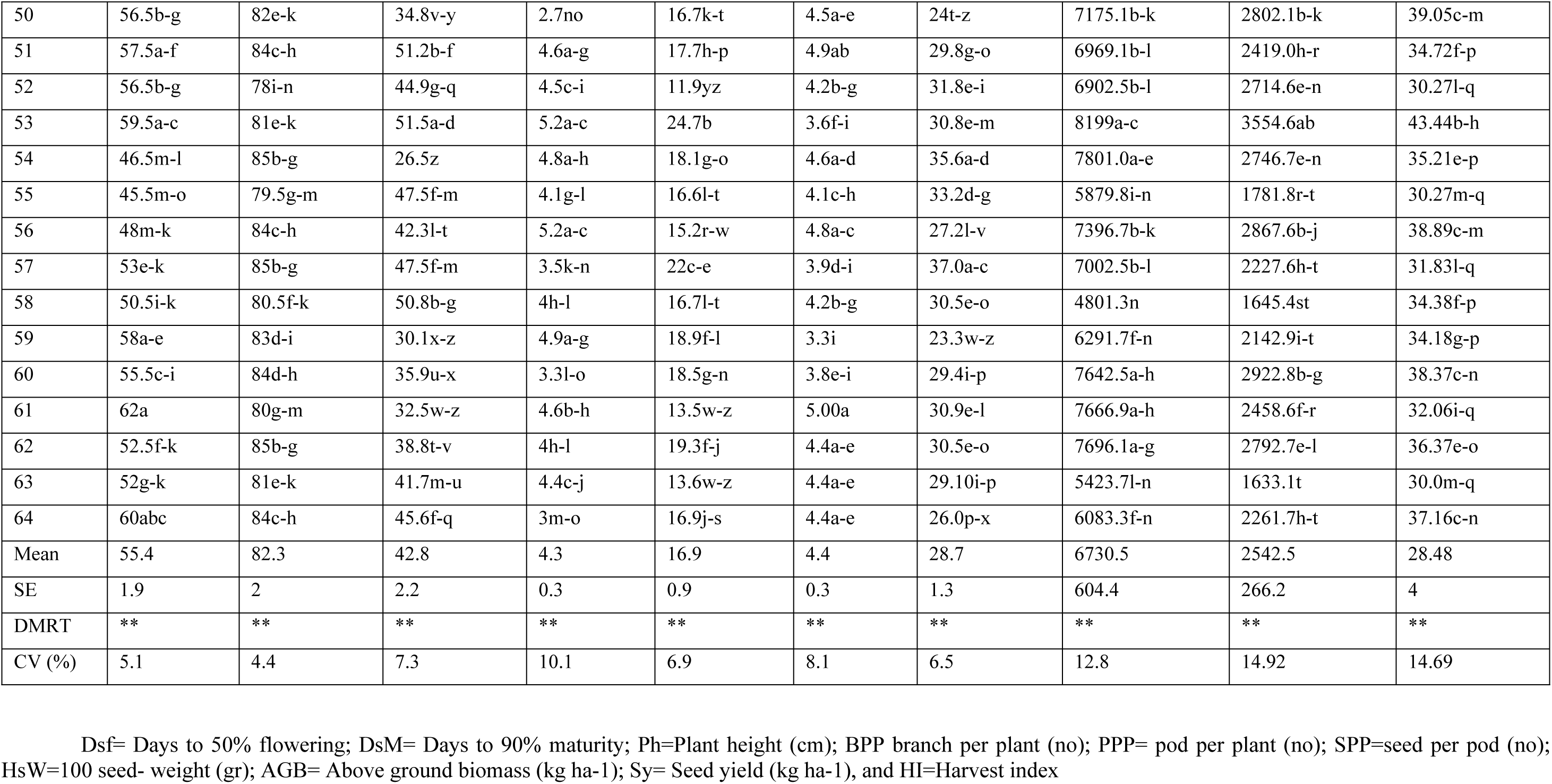
Mean values of 64 white small seeded common bean genotypes.

